# Loss of TBK1 activity leads to TDP-43 proteinopathy through lysosomal dysfunction in human motor neurons

**DOI:** 10.1101/2021.10.11.464011

**Authors:** Jin Hao, Michael F. Wells, Gengle Niu, Irune Guerra San Juan, Francesco Limone, Atsushi Fukuda, Marcel F Leyton-Jaimes, Brian Joseph, Menglu Qian, Daniel A. Mordes, Bogdan Budnik, Zhixun Dou, Kevin Eggan

## Abstract

Amyotrophic lateral sclerosis (ALS) is a fatal neurodegenerative disease characterized by motor neuron loss accompanied by cytoplasmic localization of TDP-43 proteins and their insoluble accumulations. Haploinsufficiency of TBK1 has been found to associate with or cause ALS. However, the cell-autonomous mechanisms by which reduced TBK1 activity contributes to human motor neuron pathology remain elusive. Here, we generated a human cellular model harboring loss-of-function mutations of *TBK1* by gene editing and found that TBK1 deficiency was sufficient to cause TDP-43 pathology in human motor neurons. In addition to its functions in autophagy, we found that TBK1 interacted with endosomes and was required for normal endosomal maturation and subsequent lysosomal acidification. Surprisingly, TDP-43 pathology resulted more from the dysfunctional endo-lysosomal pathway than the previously recognized autophagy inhibition mechanism. Restoring TBK1 levels ameliorated lysosomal dysfunction and TDP-43 pathology and maintained normal motor neuron homeostasis. Notably, using patient-derived motor neurons, we found that haploinsufficiency of TBK1 sensitized neurons to lysosomal stress, and chemical regulators of endosomal maturation rescued the neurodegenerative process. Together, our results revealed the mechanism of TBK1 in maintaining TDP-43 and motor neuron homeostasis and suggested that modulating endosomal maturation was able to rescue neurodegenerative disease phenotypes caused by TBK1 deficiency.

## Introduction

Amyotrophic lateral sclerosis (ALS) is a neurodegenerative disease characterized by progressive degeneration of motor neurons (MNs), resulting in paralysis and death (Rowland and Shneider, 2001). About 10% of ALS cases are inherited from families, which is almost always a dominant trait with high penetrance (Brown and Al-Chalabi, 2017). Recently, next-generation sequencing techniques have discovered that loss-of-function mutations in the gene encoding *TBK1* are associated with neurodegenerative disorders of ALS and frontotemporal dementia (FTD) (Cirulli et al., 2015; Freischmidt et al., 2015). Additional *TBK1* (OMIM 604834) variants have also been found in patients with ALS or FTD, making *TBK1* mutations the third or fourth most frequent cause of the diseases in the studied populations (Gijselinck et al., 2015; Le Ber et al., 2015; Pottier et al., 2015; Tsai et al., 2016; Williams et al., 2015). Like many other ALS and FTD-associated genes, *TBK1* variants have been related to the full phenotypic spectrum of ALS and FTD syndromes even within the same family (Freischmidt et al., 2015; Gijselinck et al., 2015).

TBK1 is an IKK family kinase previously identified for a central role in controlling type-I interferon production in response to viral infection. It is also involved in regulating multiple cellular pathways, including inflammation, immune response, cell proliferation, autophagy, and insulin signaling (Helgason et al., 2013). Although TBK1 is a multifunctional kinase involved in many biological pathways, determining which specific downstream functions of TBK1 contribute to the development of ALS and FTD remains unresolved. A previous study has shown that the reduced *Tbk1* expression synergizes with aging mediated RIPK1 inhibition to promote neuroinflammation and neurodegeneration in mouse models (Xu et al., 2018). In this study, heterozygous *Tbk1* mutations might function as a genetic risk factor that predisposes neuroinflammation and neurodegeneration under specific environmental or physiological stresses. Additional studies also reported that loss-of-function mutations of *Tbk1* in mouse motor neurons do not display any neurodegeneration phenotype, but TBK1 might contribute to the onset of disease in SOD1^G93A^ mouse models (Brenner et al., 2019; Gerbino et al., 2020). These studies suggested that loss of TBK1 activity contributed to mouse motor neuron pathology but might affect disease progression in a non-cell-autonomous manner. A recent study using iPSC-derived TBK1 patient human motor neurons (hMNs) showed decreased cell viability due to disrupted autophagy pathways (Catanese et al., 2019), raising the question of whether cell-autonomous contributions of TBK1 in human motor neurons, which are known to degenerate in ALS patients, are central to disease pathology.

Resolving the cell-autonomous functions of TBK1 in hMNs remains critical because it could provide insights into whether the reduction of TBK1 protein found in patients directly contribute to motor neuron pathology. Furthermore, efforts are being made to develop therapeutics targeting autophagy (Nguyen et al., 2019), which is currently considered a central disease mechanism for TBK1 related ALS and FTD. Therefore, determine whether loss of autophagy activity is the crucial disease-causing mechanism for *TBK1* loss-of-function mutations specifically in hMNs could be therapeutically beneficial. In addition, TDP-43 pathology has been observed in the spinal cord of patients with *TBK1* mutations (Van Mossevelde et al., 2016), but not in *Tbk1* mutant mice. It will be necessary to systematically evaluate the effects of loss-of-function mutations of *TBK1* in human cell models.

Here, we generated human embryonic stem cells (ESCs) harboring loss-of-function mutations of *TBK1* by CRISPR/Cas9. We found that *TBK1*^−/−^ cells showed impaired autophagic flux in human cellular models and increased cytoplasmic and insoluble TDP-43 in TBK1^−/−^ hMNs. Functional analysis using MEA and axotomy also validated the roles of TBK1 in early-stage disease progression. Notably, blocking of autophagy initiation by ATG7 shRNA hairpin failed to reproduce TDP-43 pathology found in TBK1^−/−^ hMNs, suggesting a mechanism independent of autophagy. Using phosphoproteomics approach, we identified abnormal phosphorylation activities in the endosomal transport pathway in TBK1^−/−^ hMNs. We validated that loss of TBK1 activity led to impaired endosomal maturation and subsequent lysosomal dysfunction. Lysosomal inhibition sensitized TBK1^−/−^ hMNs and TBK1 patient-derived hMNs to TDP-43 pathology and neurodegeneration. Restoration of TBK1 protein expression in the loss-of-function cells restored lysosomal functions and attenuated TDP-43 pathology and neurodegenerative phenotypes. Furthermore, small molecules targeting endo-lysosomal pathway rescued neurodegenerative disease phenotypes in TBK1 patient-derived hMNs. Therefore, we conclude that loss-of-function mutations of *TBK1* are sufficient to cause TDP-43 pathology and early-stage neurodegenerative phenotypes in hMNs, through endo-lysosomal pathways. Restoration of the functional TBK1 expression and small molecules targeting the endo-lysosomal pathway might be therapeutic targets for ALS and FTD patients with *TBK1* mutations.

## Results

### Loss of TBK1 activity diminishes autophagic flux in human cellular models

To investigate whether the reduced TBK1 protein expression is relevant to ALS pathology, we used CRISPR/Cas9-mediated genome editing to introduce mutations in the human embryonic stem cell (hESC) line HUES3 *Hb9*::GFP (Figure 1A) (Di Giorgio et al., 2008). To generate the loss-of-function mutations without introducing a large truncation in the genome, we designed sgRNAs targeting the CCD1 domain of TBK1 (Figure 1A), which had been reported to cause the instability of TBK1 protein (Ye et al., 2019). After isolating and expanding single clones, two potential TBK1^−/−^ clones were validated in cells by Sanger sequencing (Figure 1B, S1A, and S1B). Western blot and immunocytochemistry (ICC) analysis confirmed the loss of TBK1 expression, as well as the increased level of SQSTM1/P62 in the two candidate clones (p<0.01) (Figure 1C, 1D, and S1C). The elevated P62 level indicated impaired autophagic flux, as TBK1 could mediate autophagic flux and cargo selection by phosphorylating autophagic receptors, like P62. For clarification, we used candidate 1, which has a one bp deletion at the desired mutant location, for future experiments. We next tested how the loss of TBK1 activity affected autophagy and confirmed the role of TBK1 in autophagosomal maturation as measured by LC3 immunoblotting (Figure 1E and 1F). While control cells showed increased LC3-II (an early autophagic organelle marker) upon lysosomal inhibitor bafilomycin A1 (Baf A1) treatment, TBK1^−/−^ hESCs accumulated LC3-II in the basal level but failed to further increase upon Baf A1 treatment. This indicates a deficiency in autophagosome maturation in the TBK1^−/−^ cells, consistent with a previous report (Pilli et al., 2012).

**Figure 1.**
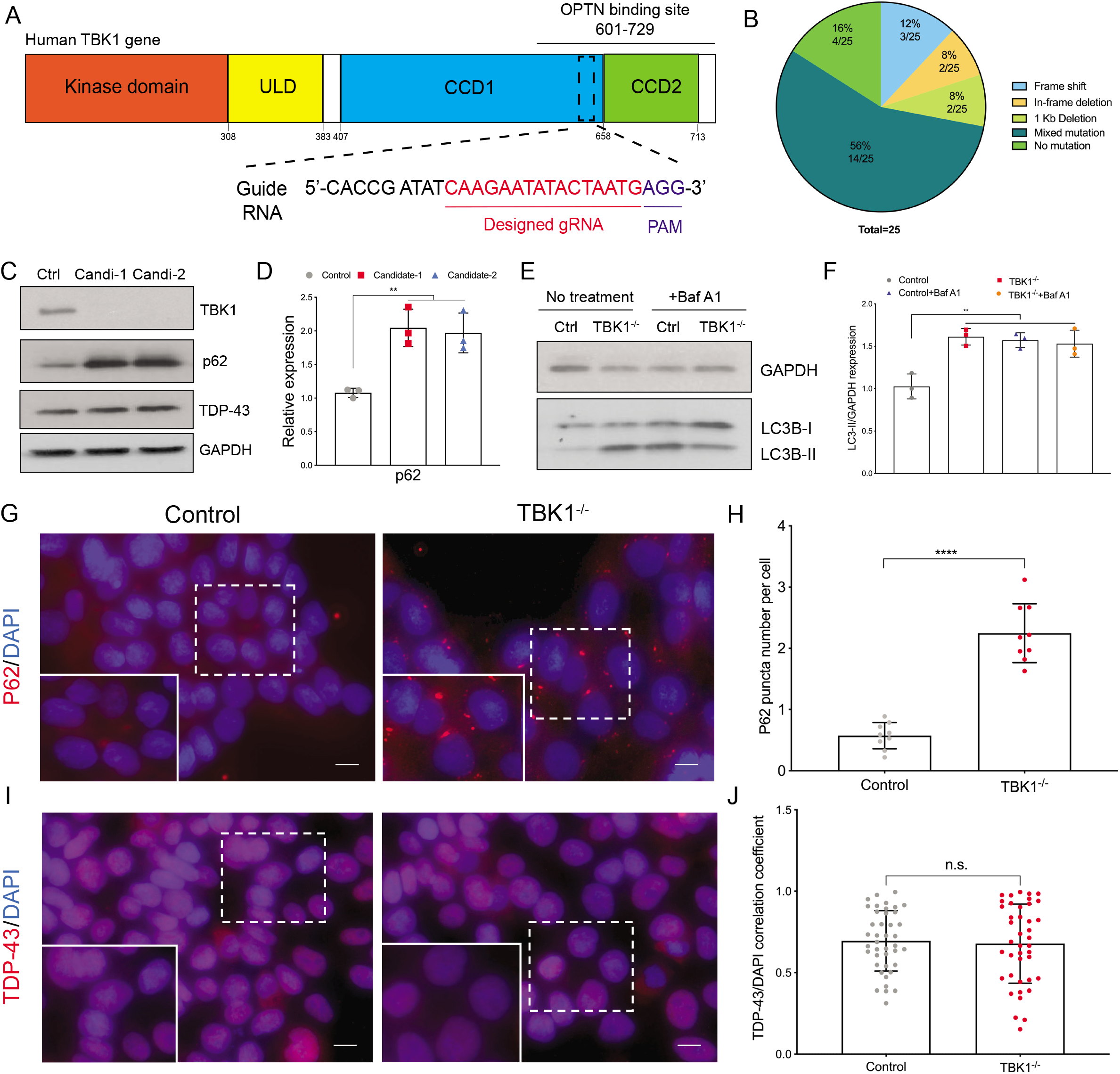
Loss-of-function mutations of TBK1 cause deficient autophagic flux. (A) Schematic of gene-editing strategy and targeting locus of sgRNA for human TBK1 gene in HUES3 *HB9*::GFP ESCs. (B) Sanger sequencing results of selected colonies for TBK1 mutations at the indicated sites. (C) Validation of TBK1, P62, and TDP-43 protein expression for two candidate mutant clones by western blot. (D) Quantifications of the protein levels in (C) from 3 independent experiments are shown. (E) Effects of TBK-1 on LC3-II levels and degradation during autophagic maturation. Control and TBK1^−/−^ cells are treated or not treated with bafilomycin A1 (BafA1) to inhibit autophagic degradation of LC3-II. (F) Quantification of LC3-II level is shown from 3 independent experiments. Levels of LC3-II expression are normalized to GAPDH. (G-J) Immunostaining of P62 (G) and TDP-43 (I) for hESC cultures of control and TBK1^−/−^ cells. Scale bar, 20 μm. Quantifications of p62 puncta numbers (H) and TDP-43/DAPI correlation coefficient (J) are shown. Data represent means ± SE (n ≥ 3; ns means p ≥ 0.05; **p < 0.01; ****p < 0.0001; student’s t-test and ANOVA).

Next, we sought to determine whether the clearance of P62 puncta, which has been reported to be an ALS pathological hallmark, would be affected by TBK1 deficiency. We performed ICC and quantitative imaging analysis and detected significantly increased P62 puncta in TBK1^−/−^ cells (p<0.0001) (Figure 1G and 1H), consistent with previous protein blot analysis (Figure 1C and 1D). Therefore, P62 puncta cannot be cleared when TBK1 is deficient. In addition, the accumulated P62 puncta at the peri-nuclear sites indicated blocked autophagic degradative pathways, which has been reported in *TBK1* mutant and other sporadic ALS patients (Mizuno et al., 2006; Van Mossevelde et al., 2016). Next, we sought to determine whether loss of TBK1 activity leads to mislocalization of TDP-43, another critical pathological hallmark for ALS, in hESC models. We deployed immunocytochemistry analysis using endogenous TDP-43 antibody (Figure 1I) and compared the correlation coefficient of TDP-43 with nuclear staining DAPI (Figure 1J). However, we detected no mislocalized TDP-43 in TBK1^−/−^ cells compared to the control cells (p=0.7161). Protein blot analysis using subcellular fractionation also confirmed that TBK1^−/−^ cells did not show any aberrant TDP-43 localizations (Figure S1D). Thus, we generated a human stem cell model with loss-of-function mutations of *TBK1*, and loss of TBK1 activity induced P62, but not TDP-43 pathology, in human stem cell models.

### TBK1 ablation leads to TDP-43 pathology in hMNs

To investigate the functional consequences of TBK1 mutation in hMNs, the most ALS-relevant cell type, we differentiated the control and TBK1^−/−^ hESCs transfected by NGN2 into motor neurons using the NGN2 overexpression method adapted from previous protocols (Figure S2A) (Nehme et al., 2018). Over 90% of cells showed *Hb9*::GFP positive signals during motor neuron patterning (Figure S2B). To determine the efficiency of motor neuron differentiation in both control and TBK1^−/−^ cells, we used ICC to quantify the percentage of ISL-1 positive cells and found that TBK1 deficiency did not affect the differentiation efficiency (p=0.9757) (Figure S2D).

To determine whether loss of TBK1 activity in hMNs had a similar effect on key ALS protein expression as in hESCs, we first performed protein blot analysis to confirm the increased expression of P62 (p<0.0001) and TDP-43 (candidate-1: p=0.9987, candidate-2: p=0.7270) in hMNs, which is consistent with the previous result in hESCs (Fig S2C). Although TDP-43 pathology was not detected in TBK1^−/−^ hESCs, it is intriguing to investigate this effect in hMNs, which has been reported to display TDP-43 proteinopathies in ALS patients with reduced TBK1 expressions (Van Mossevelde et al., 2016). To determine whether TBK1 activity affects TDP-43 homeostasis in hMNs, we performed ICC and immunoblot analysis using differentiated hMNs on day 3 after plating (Figure 2A). Interestingly, TBK1^−/−^ hMNs led to increased cytoplasmic localization of TDP-43, a key feature of TDP-43 proteinopathies in ALS/FTD (Figure 2B). The correlation coefficient analysis of TDP-43 with nuclear staining DAPI confirmed the decreased overlapping of TDP-43 with the nucleus in TBK1^−/−^ hMNs (p<0.0001) (Figure 2C). To further determine the subcellular localization of TDP-43 in TBK1^−/−^ hMNs, we performed subcellular fractionation followed by immunoblotting (Figure 2D). In the control hMNs, TDP-43 was primarily localized in the nucleus, while loss of TBK1 activity led to a dominant localization of TDP-43 in the pellet fraction (Figure 2E and quantified in 2F). Our results indicate that loss of TBK1 activity is sufficient to cause cytoplasmic TDP-43 and their insoluble accumulations in hMNs.

**Figure 2.**
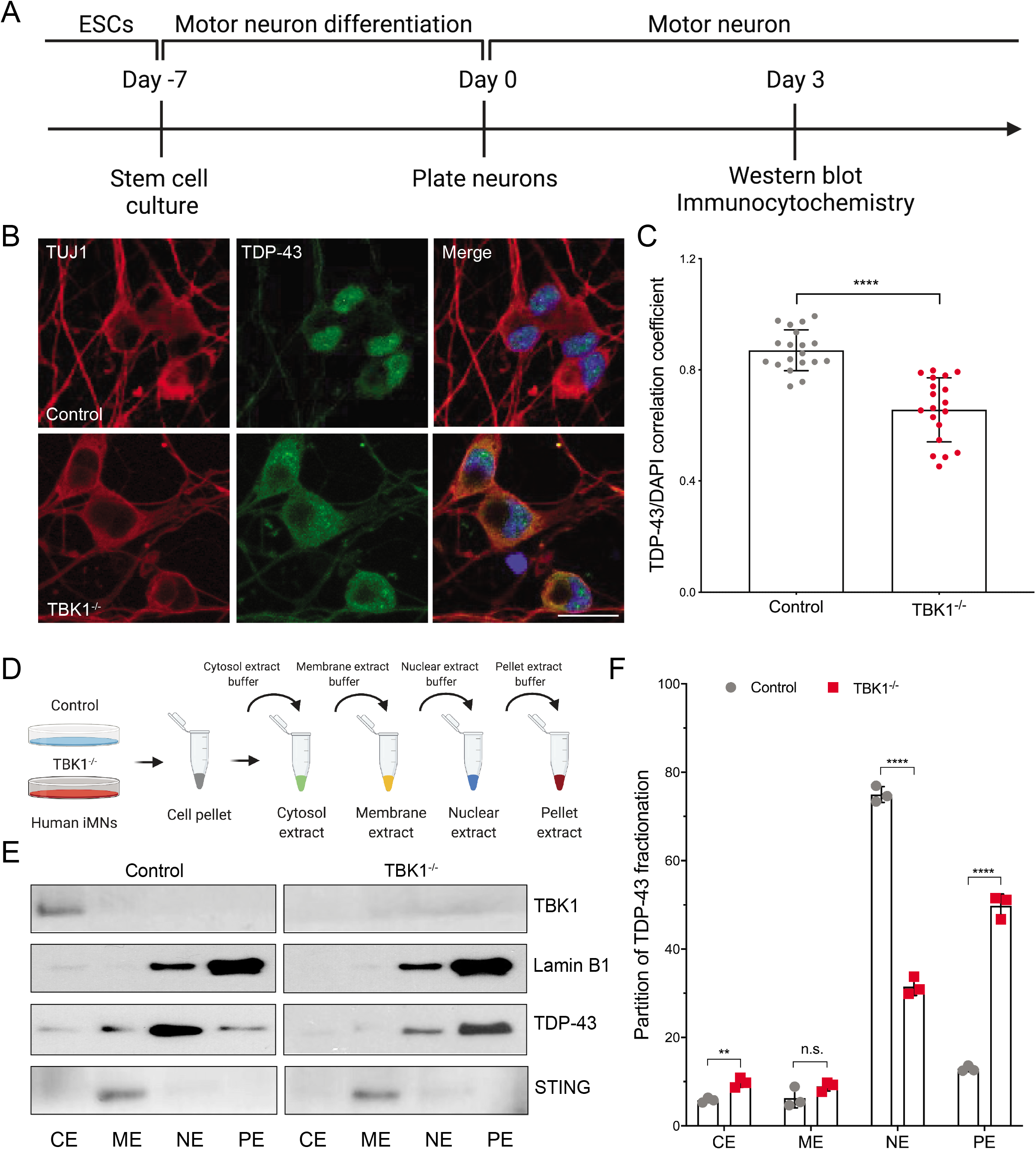
Loss of TBK1 activity leads to TDP-43 pathology in human motor neurons. (A) Experimental outline of TDP-43 localization assay in hMNs. Motor neurons are differentiated, and TDP-43 localization is quantified by immunocytochemistry on Day 3 after replating. Subcellular fractionation of TDP-43 is quantified by western blot in Day 3 hMNs. (B) Representative images of Day 3 motor neurons for quantification of TDP-43 localization. All cells are stained by DAPI, TDP-43, and neuronal marker TUJ1. Scale bar, 20 μm. (C) Quantifications of the TDP-43/DAPI correlation coefficient from (B) are shown. (D) Schematic of subcellular fractionation experiment for control and TBK1^−/−^ hMNs. Sequential extracts of cytosol, membrane, nuclear, and pellet fractions are performed. (E) Subcellular fractionation analysis of TBK1 (cytosol control), Lamin B1 (nuclear and pellet control), STING (membrane control), and TDP-43 levels in different fractions of control and TBK1^−/−^ hMNs. Representative blots show TDP-43 levels in cytosol extract (CE), membrane extract (ME), nuclear extract (NE), and pellet extract (PE). (F) Quantifications of the partition of TDP-43 in each cellular fraction for control and TBK1^−/−^ hMNs are shown. Data represent means ± SE (n ≥ 3; ns means p ≥ 0.05; **p < 0.01; ***p < 0.001; ****p < 0.0001; student’s t-test and ANOVA).

### Autophagy inhibition fails to induce TDP-43 pathology in hMNs

Loss-of-function mutations of TBK1 caused impaired autophagic flux, and the autophagy pathways were considered a key mechanism for ALS progression. To investigate if loss of TBK1 contributed to pathological TDP-43 accumulations through the autophagy pathway, we compromised autophagy and measured the amount of insoluble TDP-43 accumulation in hMNs. Specifically, we treated motor neurons with DMSO (vehicle control) or 100 nM Baf A1 for 24 hr (Figure 3A). The treatment of Baf A1 did not significantly affect the amount of soluble and insoluble TDP-43 in control motor neurons (Figure 3B and 3C), indicating that insoluble TDP-43 accumulation was not initiated by autophagy inhibition. By contrast, TBK1^−/−^ motor neurons accumulated about 2-fold more insoluble TDP-43 (p<0.0001), while soluble TDP-43 level was reduced by about 50% when Baf A1 was added (p<0.05) (Figure 3B and 3C). In addition, when we compromised the proteasome activity proteasome inhibitor MG132, fractions of soluble TDP-43 were reduced in control hMNs (p=0.03), while insoluble forms of TDP-43 were not substantially altered in both control and TBK1^−/−^ motor neurons (p>0.99) (Figure S3A).

**Figure 3.**
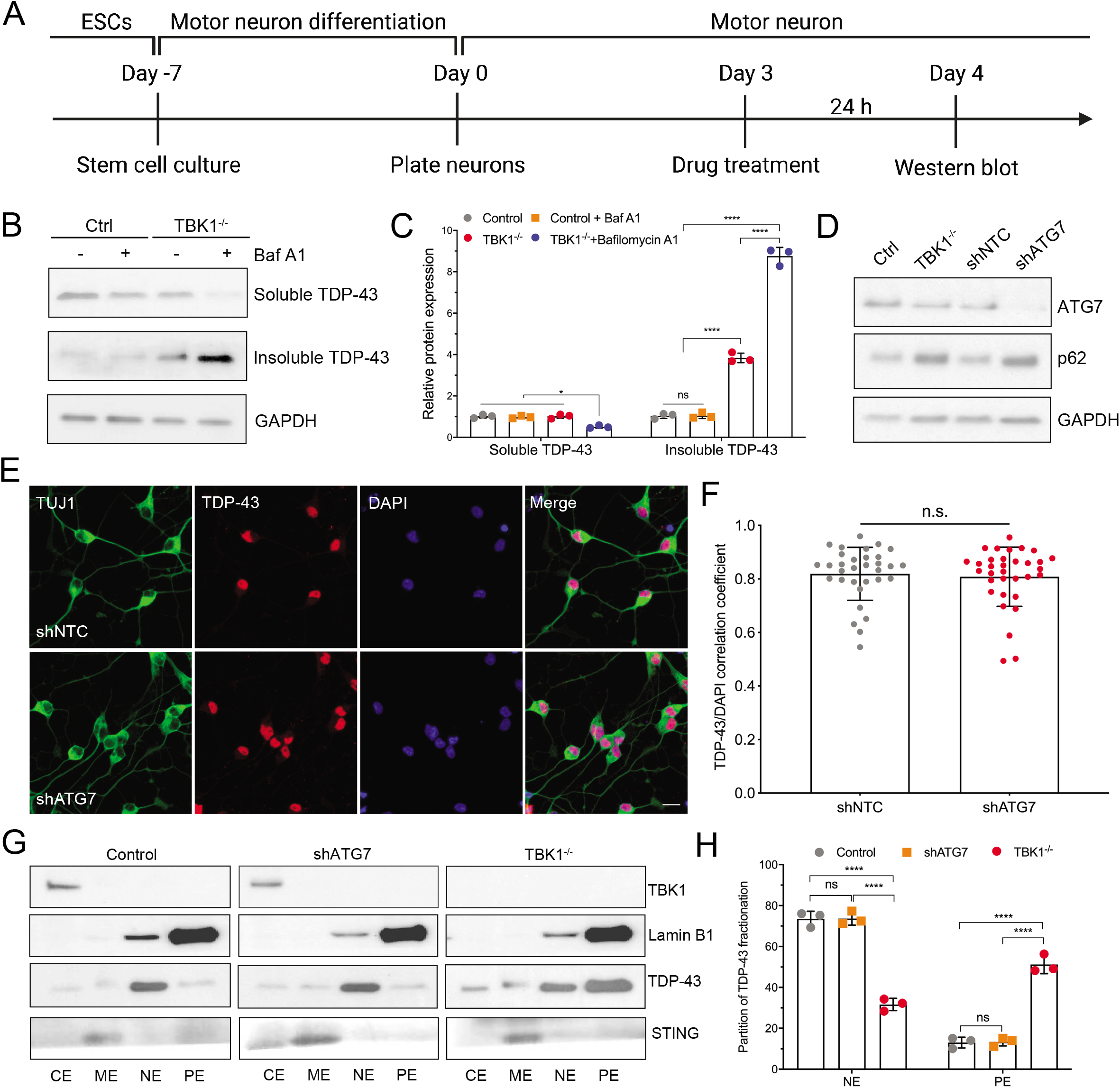
Insoluble TDP-43 accumulation induced by TBK1^−/−^ is independent of autophagy activity. (A) Experimental outline of western blot analysis for insoluble TDP-43. Control and TBK1^−/−^ motor neurons are treated with DMSO or 100 nM Baf A1 for 24 hr on Day 3. Western blot analysis is performed on Day 4. (B) Western blot analysis of control and TBK1^−/−^ hMNs treated with DMSO or Baf A1. Accumulations of TDP-43 proteins are shown in the soluble (RIPA) and insoluble (urea) fractions. (C) Quantification of TDP-43 expression between DMSO and Baf A1 treated hMNs. GAPDH in the soluble fraction is used as the loading control for normalization. (D) Western blot analysis of hMNs stably expressing non-targeting control (shNTC) or shATG7 hairpin. The expression of ATG7 and p62 is normalized to GAPDH (loading control). (E) Representative images of shNTC and shATG7 motor neurons for quantification of TDP-43 localization. All cells are stained by DAPI, TDP-43, and neuronal marker TUJ1. Scale bar, 20 μm. (F) Quantifications of the TDP-43/DAPI correlation coefficient from (E) are shown. (G) Subcellular fractionation analysis of TBK1 (cytosol control), Lamin B1 (nuclear and pellet control), STING (membrane control), and TDP-43 levels in different fractions of control, TBK1^−/−^, and shATG7 hMNs. (H) Quantifications of the partition of TDP-43 in each cellular fraction for control, TBK1^−/−^, and shATG7 hMNs are shown. Data represent means ± SE (n ≥ 3; ns means p ≥ 0.05; *p < 0.05; ****p < 0.0001; student’s t-test and ANOVA).

Next, to evaluate if the increased level of insoluble TDP-43 affected the survival of control and TBK1^−/−^ motor neurons, Day 2 motor neurons were treated with different concentrations of MG132 or Baf A1 for 48 hr for subsequent survival assay (Figure S3B). Compared to control motor neurons, treatment of MG132 did not affect the survival of TBK1^−/−^ motor neurons, while Baf A1 treated TBK1^−/−^ motor neurons showed significantly reduced survival in a variety of concentrations (p<0.0001) (Figure S3C). Therefore, the treatment of Baf A1 could prevent TBK1^−/−^ motor neurons from degrading pathological TDP-43 proteins, increasing insoluble TDP-43 accumulations, and hindering hMNs survival to treatment.

Considering the multifaced function of Baf A1, which could block autophagosome-lysosome fusion and inhibit endosomal acidification, we investigated the role of selective autophagy inhibition in regulating TDP-43 homeostasis by shRNA knockdown of ATG7. We generated cells stably expressing non-targeting control (shNTC) and ATG7 shRNA (shATG7) hairpin. Immunoblotting confirmed that ATG7 was deficient in shATG7 hMNs, and ATG7 deficiency caused increased levels of P62 and impaired autophagic flux (Figure 3D and S3D). After motor neuron differentiation, the shATG7 did not affect the percentage of cells expressing motor neuron marker, ISL-1, compared to the shNTC control (p=0.4417) (Figure S3E and S3F). Interestingly, we found that inhibition of ATG7 did not alter the localization of TDP-43, as it stayed mainly in the nucleus (Figure 3E and 3F). To determine if ATG7 inhibition affected the localization of TDP-43 expression, we performed subcellular fractionation and immunoblotting. We found that ATG7-deficient hMNs showed similar levels of TDP-43 in different fractions compared to control hMNs (NE: p=0.9983, PE: p=0.9885) (Figure 3G and 3H). These data suggest that defective autophagy is unlikely to be the major cause for TDP-43 pathology in hMNs, and TBK1 deficiency might initiate TDP-43 pathology through autophagy-independent pathways.

### TBK1 deficiency impairs endosomal maturation and lysosomal function

Determining the functional pathway in which TBK1 acts may be beneficial for designing therapeutics treating ALS. To better understand the mechanisms of TBK1^−/−^ induced TDP-43 pathology and neurodegeneration, we performed phosphoproteome analysis for control, shATG7, and TBK1^−/−^ motor neurons using TMT labeling (Figure 4A). Compared to control motor neurons, TBK1^−/−^, but not shATG7 motor neurons, showed markedly reduced phosphorylation levels of proteins involved in nuclear transport and endosomal transport signaling pathways (Figure 4B and 4C). Previous reports also showed that TBK1 was required for early endosomal targeting, and TBK1 might modulate RAB GTPase activity (Fraser et al., 2019; Heo et al., 2018; Pilli et al., 2012). To determine whether loss of TBK1 caused changes in endosomal transport in hMNs, we evaluated the expression of markers for early endosomes (EEA1, RAB5) and late endosomes (RAB7) in control and TBK1^−/−^ motor neurons by ICC. We identified an increased number of EEA1-positive and RAB5-positive vesicles but reduced RAB7-positive vesicles in TBK1^−/−^ motor neurons using confocal microscopy (p<0.0001) (Figure 4D-G). Our results suggested that TBK1 might mediate the maturation of early endosomes to late endosomes, and reduced RAB7 level caused by loss of TBK1 might indicate the impaired maturation and fusion of late endosomes with lysosomes (Figure S4A).

**Figure 4.**
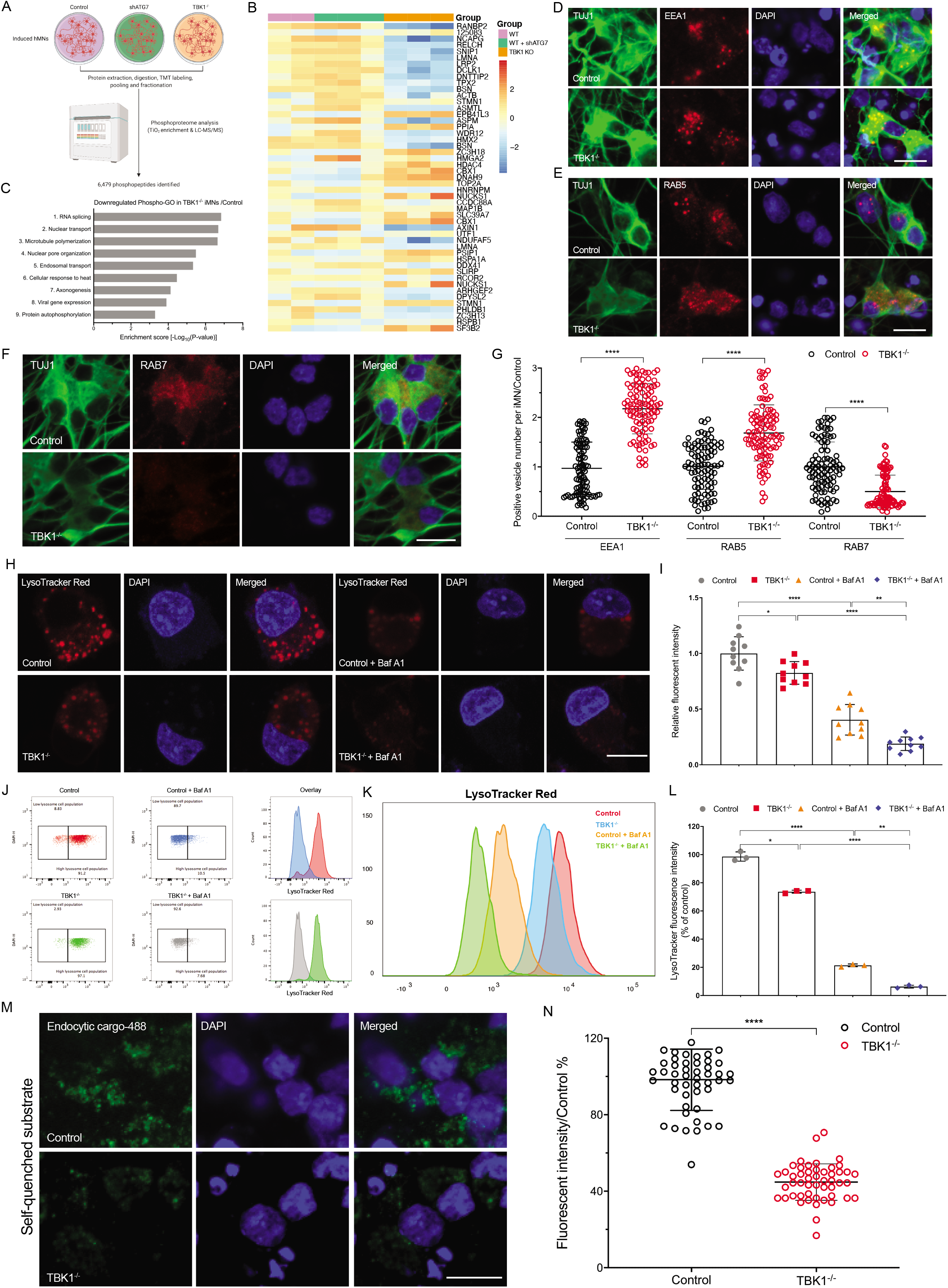
Loss of TBK1 activity leads to impaired endosomal maturation and defective lysosomal function. (A) Schematic of experimental design for phospho-proteomic analysis from control, TBK1^−/−^, and shATG7 hMNs. (B) Heatmap of the 50 most significant differentially expressed phospho-proteins (log2 expression). (C) GO analysis of proteins that are down phosphorylated in TBK1^−/−^ hMNs compared to control hMNs. (D-G) Representative images of control and TBK1^−/−^ motor neurons for vesicle number quantification of EEA1 (D), RAB5 (E), and RAB7 (F). All cells are stained by DAPI and neuronal marker TUJ1. Scale bar, 20 μm. Quantifications of the positive vesicle numbers in TBK1^−/−^ hMNs from (D-F) are shown. All quantifications are normalized to control hMNs (G). (H) LysoTracker Red staining of control and TBK1^−/−^ hMNs. The control cells show big bright puncta, while the TBK1^−/−^ cells have much dimmer and fewer puncta. Baf A1 is used as a positive control as the acidification inhibitor. (I) Quantification of LysoTracker relative intensity from (H) is shown. (J-L) Flow cytometry analysis of LysoTracker Red staining in control and TBK1^−/−^ hMNs. Baf A1 is added as a positive control and shows decreased LysoTracker Red staining (J). The intensity of LysoTracker Red in control, TBK1^−/−^, control + Baf A1, and TBK1^−/−^ + Baf A1 hMNs are compared (K). Quantifications of LysoTracker Red fluorescent intensity are shown (L). (M) Lysosome enzyme activity assay in control and TBK1^−/−^ hMNs. Self-quenched substrate is used as endocytic cargo, and fluorescence signal, generated by lysosomal degradation, is proportional to the intracellular lysosomal activity. (N) Quantifications of the relative fluorescent intensity from (M) are shown. Data represent means ± SE (n ≥ 3; ns means p ≥ 0.05; *p < 0.05; ***p < 0.001; ****p < 0.0001; ANOVA and student’s t-test).

To determine the functional consequences of impaired endosomal pathway and its link with TDP-43 pathology, we sought to examine the lysosomal function, which is downstream of endosomal pathway, in TBK1^−/−^ hMNs. Our study demonstrated that TBK1 mediates the maturation of autophagosomes and endosomes, thereby prevents their fusion with lysosomes. Previous report shows that the functional activation of lysosomes is regulated by the lysosomal fusion process (Zhou et al., 2013). To determine if loss of TBK1 affected lysosomal biogenesis, we analyzed the the number of LAMP1-positive vesicles, in control and TBK1^−/−^ motor neurons by ICC (Figure S4B and S4C). TBK1^−/−^ motor neurons showed an increased number of LAMP1-positive vesicles than control motor neurons (p<0.0001), indicating that TBK1 deletion in motor neurons might lead to accumulation of lysosomes. The other critical function of lysosomes is being acidic, which can be mediated by degradation of lysosomal-substrates (Hu et al., 2015). To assess whether these LAMP1-positive vesicles were acidic, a functional indicator of lysosomes, motor neurons were labeled with LysoTracker red, an acidic organelle staining dye. Surprisingly, we found that LysoTracker-positive vesicles were reduced in TBK1^−/−^ motor neurons (p<0.0001), and Baf A1, a lysosomal acidification inhibitor, diminished the LysoTracker staining in motor neurons (Figure 4H and 4I). Next, we quantified the LysoTracker Red fluorescence intensity in motor neurons using flow cytometry. Consistent with LysoTracker staining data, total LysoTracker intensity was markedly reduced in TBK1^−/−^ motor neurons (p<0.05) (Figure 4J-L), suggesting that lysosomes were less acidic in TBK1^−/−^ motor neurons. Subsequently, we used the self-quenched enzymatic substrates, which would be targeted to the lysosomes by endocytosis and turn fluorescent upon the enzymatic activity to quantify endo-lysosomal activities (Humphries and Payne, 2012). Consistent with the impaired endosomal maturation and deficient lysosomal enzyme activities observed in TBK1^−/−^ motor neurons, fluorescent endocytic cargos upon endo-lysosomal transport and lysosomal enzymatic activity were reduced by ~60% relative to control cells (p<0.0001) (Figure 4M and 4N). Together, our results suggest that loss of TBK1 impairs the endosomal maturation and leads to deficient lysosomal enzymatic activities with reduced acidifications.

### Inhibiting lysosomal acidification triggers TDP-43 pathology in hMNs

Our data suggests that TBK1^−/−^ hMNs accumulates cytoplasmic and insoluble TDP-43 as well as impaired endosomal maturation and subsequent lysosomal dysfunction. To establish the link between TDP-43 pathology and the lysosomes, motor neurons were treated with Lys05, which blocks lysosomal enzymatic activity by neutralizing lysosomal pH (Amaravadi and Winkler, 2012) (Figure 5A). To validate the role of TBK1 in endo-lysosomal pathway, we co-stained active phosphorylated TBK1 (pTBK1) with key markers of early endosomes under lysosomal stress. Consistent with a previous study (Fraser et al., 2019), pTBK1 stained positively with EEA1 vesicles when lysosomal activity was inhibited (Figure 5B and 5C), suggesting that TBK1 activity lies upstream of endo-lysosomal pathways. The fact that pTBK1 co-localize with early endosomal marker, EEA1, is consistent with our aforementioned data that loss of TBK1 caused deficient lysosomal function and suggests a potential link between TDP-43 pathology and the endo-lysosomal pathway.

**Figure 5.**
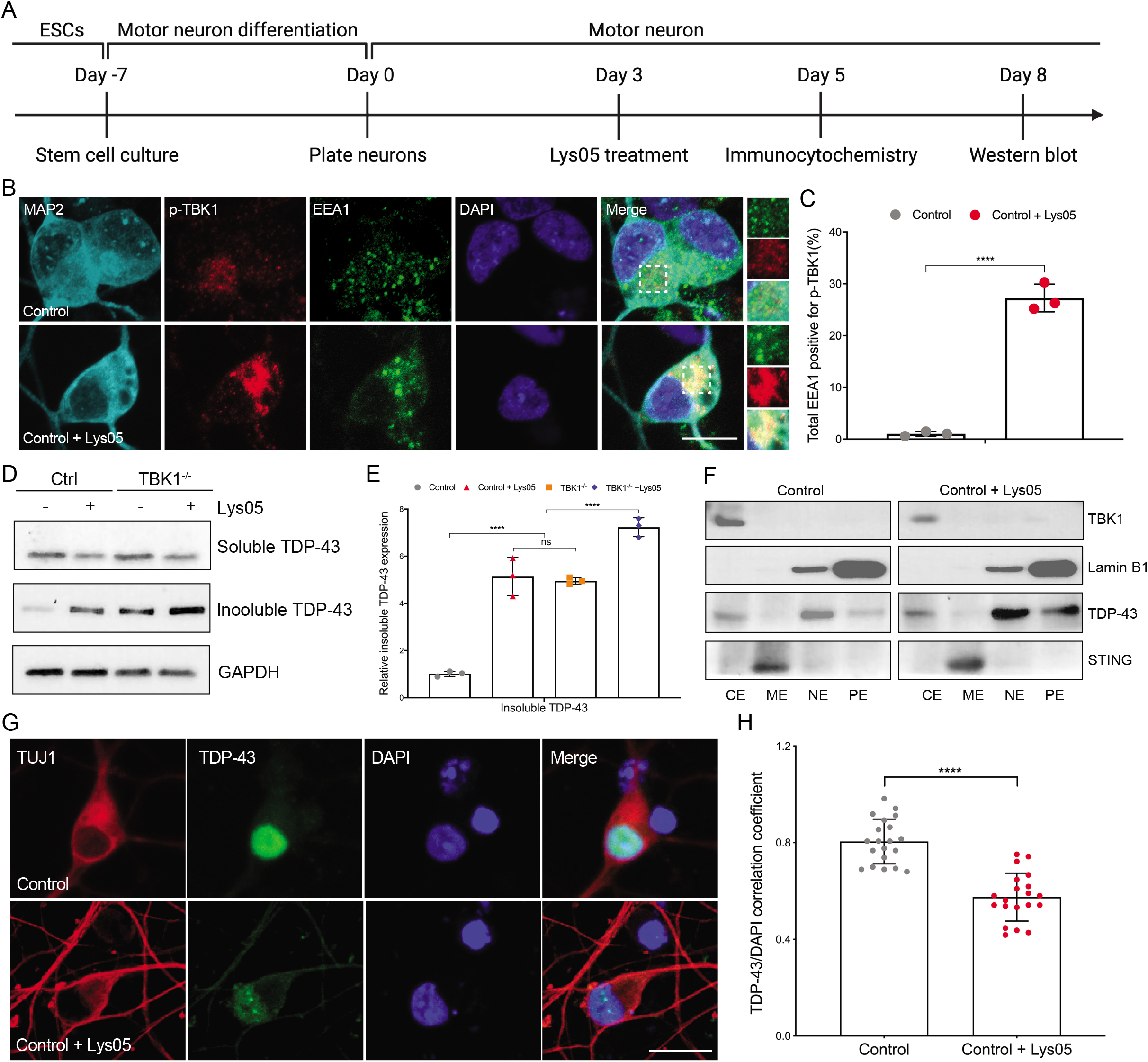
Lysosomal function deficiency is sufficient to trigger TDP-43 pathology in hMNs. (A) Schematic of experimental outline for lysosomal inhibitor treatment in hMNs. (B) Representative images of control motor neurons treated with and without Lys05 for staining against EEA1 and pTBK1. All cells are stained by DAPI, EEA1, pTBK1, and neuronal marker MAP2. Scale bar, 20 μm. (C) Quantifications of the percentage of total EEA1 vesicles showed positive for pTBK1. (D) Western blot analysis of control and TBK1^−/−^ hMNs treated with DMSO or Lys05. Accumulations of TDP-43 proteins are shown in the soluble (RIPA) and insoluble (urea) fractions. (E) Quantification of TDP-43 expression between DMSO and Lys05 treated hMNs. GAPDH in the soluble fraction is used as the loading control for normalization. (F) Subcellular fractionation analysis of TBK1 (cytosol control), Lamin B1 (nuclear and pellet control), STING (membrane control), and TDP-43 levels in different fractions of control hMNs treated with and without Lys05. Representative blots show TDP-43 levels in cytosol extract (CE), membrane extract (ME), nuclear extract (NE), and pellet extract (PE). (G) Representative images of control motor neurons treated with and without Lys05 for staining against TDP-43. All cells are stained by DAPI, TDP-43, and neuronal marker TUJ1. Scale bar, 20 μm. (H) Quantifications of the TDP-43/DAPI correlation coefficient from (G) are shown. Data represent means ± SE (n ≥ 3; ns means p ≥ 0.05; ****p < 0.0001; ANOVA and student’s t-test).

To further validate the link connecting TBK1^−/−^ induced TDP-43 pathology and the impairment of endo-lysosomal pathway, we asked whether inhibited lysosomal activity is sufficient to cause TDP-43 pathology in hMNs. To this end, we treated control and TBK1^−/−^ hMNs with 1 μM Lys05 for 4-5 days to inhibit the lysosomal activity and tested the level of insoluble TDP-43 accumulation. Lysosomal inhibition in control hMNs showed increased insoluble TDP-43 levels (p<0.0001), supporting the notion that the TBK1^−/−^ induced TDP-43 pathology might be caused by deficient lysosomal activity (Figure 5D and 5E). Interestingly, the level of insoluble TDP-43 in control hMNs treated with Lys05 was comparable to that of TBK1^−/−^ hMNs (Figure 5D and 5E), highlighting the central role of lysosomal activity in the clearance of insoluble TDP-43. Subcellular fractionation experiment also confirmed increased pellet fraction of TDP-43 in control hMNs when treated with Lys05 (Figure 5F). In addition, increased cytoplasmic TDP-43 was observed in control cells treated with Lys05 (p<0.0001) (Figure 5G and 5H), suggesting that lysosomal activity is linked to cytoplasmic TDP-43 proteins and their insoluble accumulations. Thus, our data validates the strong link between TDP-43 pathology and the activity of endo-lysosomal pathway, and TBK1 is an important mediator of these processes.

### Loss of TBK1 causes hyperexcitability and impaired axonal repair in hMNs

As loss-of-function mutations of *TBK1* caused TDP-43 proteinopathies in hMNs, we proceeded to ask if TBK1^−/−^ in hMNs could disrupt neuronal homeostatic functions. Survival of control and TBK1^−/−^ hMNs was measured on Days 14 and 28 compared to Day 2 (Figure S6A and S6B). With the culture of hMNs, about 60% of control and TBK1^−/−^ neurons survived on Day 14 (p=0.8281), and about 43% of them survived on Day 28 (p=0.9759) (Figure S6C). Therefore, loss of TBK1 activity did not affect human motor neuron survival in culture.

Although *TBK1* mutation in hMNs did not cause neuronal death, we sought to examine whether loss of TBK1 could lead to early-stage neurodegenerative phenotypes. Physiological analyses had demonstrated alterations in the functional properties of motor neurons at very early stages in transgenic mouse models of ALS. Subsequent studies had revealed pre-symptomatic hyperexcitability in ALS motor neurons (Wainger et al., 2014; Zona et al., 2006). Therefore, perturbations of the intrinsic biological properties might lead to abnormal electrophysiological activities of motor neurons, and these activities could reflect and contribute to the earliest events that caused neurodegeneration in ALS motor neurons. To determine if TBK1 ablation in hMNs caused any electrophysiological phenotype, we recorded the spontaneous firing of control and TBK1^−/−^ motor neurons using extracellular multielectrode arrays (MEAs) at Day 21 and 28 in culture (Figure 6A). We plated an equal number of *Hb9*::GFP positive control and TBK1^−/−^ motor neurons and carefully monitored their attachment on the plate throughout the experiments (Figure 6B). After four weeks of culture, the control and TBK1^−/−^ motor showed a similar number of active electrodes on MEA plates (p=0.9585) (Figure S6D). Intriguingly, TBK1^−/−^ motor neurons exhibited a significantly greater number of spikes and a higher spontaneous firing rate (p<0.0001) (Figure 6C). Furthermore, TBK1^−/−^ motor neurons consistently showed increased bursting (p<0.0001) and network bursting frequency (p<0.01) than control motor neurons (Figure 6C). Next, we sought to determine if retigabine, a specific activator of subthreshold Kv7 currents, could inhibit the spontaneous firing of TBK1^−/−^ motor neurons. Consistent with our previous report using SOD1 ALS motor neurons (Wainger et al., 2014), retigabine also stopped the spontaneous firing of TBK1^−/−^ motor neurons in a dose-dependent manner (Figure S6E).

**Figure 6.**
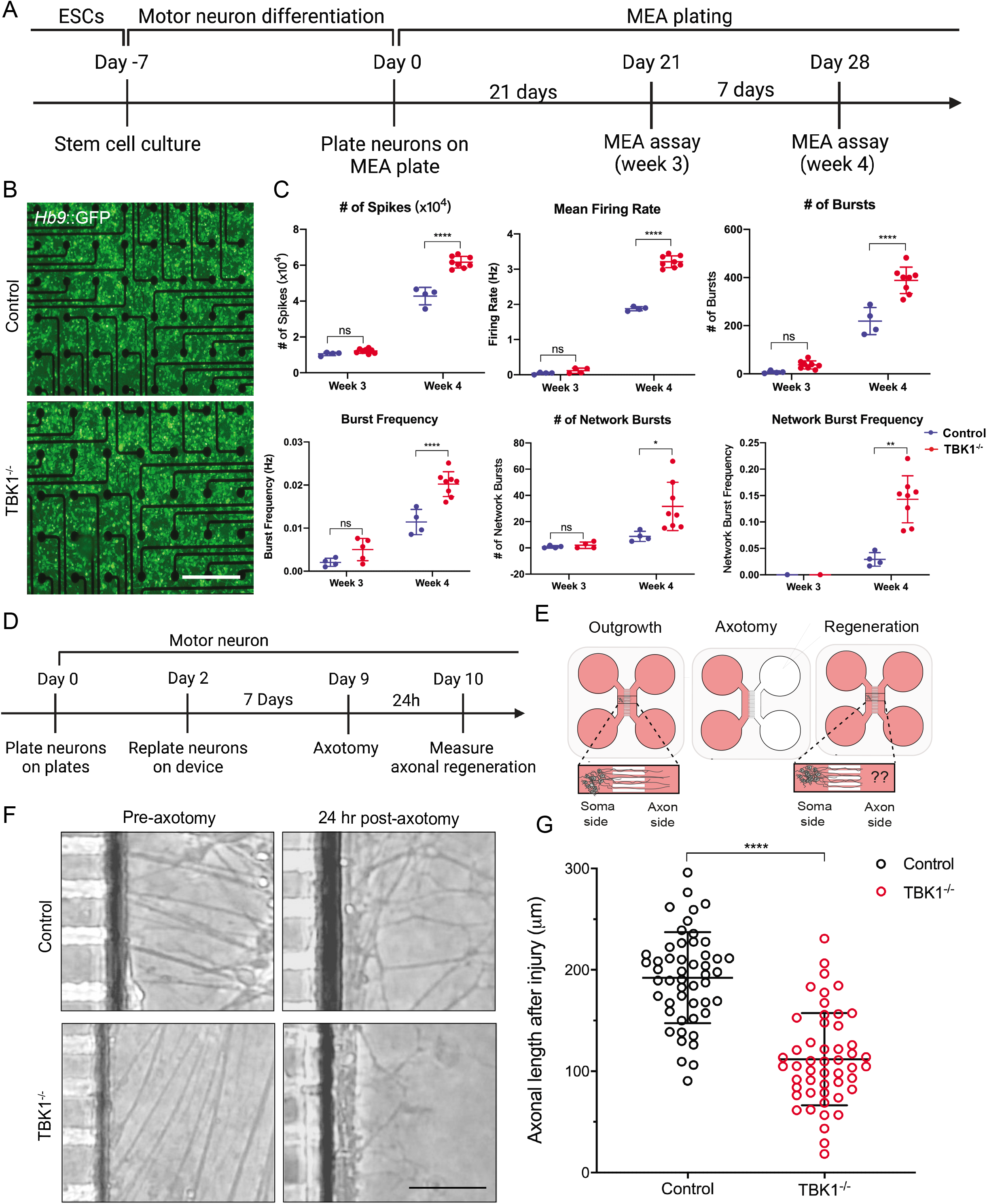
TBK1 ablation causes early-stage motor neuron degenerative phenotypes implicated in ALS. (A) Experimental outline of MEA assay in hMNs. Motor neurons are differentiated and replated on MEA plates. Electrophysiological properties of control and TBK1^−/−^ hMNs are measured on Day 21 and Day 28 after replating. (B) Representative images at ×4 magnification of *Hb9*::GFP-fluorescing control and TBK1^−/−^ motor neurons plated on MEA wells on day 14 in culture. Scale bar: 400 μm. (C) Analysis of MEA recordings from days 21 to 28 in culture. TBK1^−/−^ motor neurons show increased spike numbers and mean firing rate compared to control motor neurons, and the number of bursts, network bursts, and burst frequency are elevated. (D) Experimental outline of axotomy assay in hMNs. Control and TBK1^−/−^ hMNs are plated on the axotomy device. Axotomy is performed on Day 9, and measurement of axonal regrowth is conducted on Day 10. (E) Schematic of axonal regrowth experiment upon injury. The hMNs are plated on the left chamber (soma side) of the device, and axons sprout into the right chamber (axon side) through the channels connecting both sides. Axotomy is performed by washing out the axon side to cut the sprouted axons. The ability of axonal repair is measured by the length of re-sprouted axons. (F) Representative images of axonal sprouting in control and TBK1^−/−^ hMNs from pre- and post-axotomy conditions. Scale bar: 500 μm. (G) Quantifications of axonal regrowth length for control and TBK1^−/−^ hMNs are shown. The data of individual neurites is displayed as a single dot. Data represent means ± SE (n ≥ 3; ns means p ≥ 0.05; *p < 0.05; **p < 0.01; ***p < 0.001; ****p < 0.0001; student’s t-test and ANOVA).

In addition to hyperexcitability, distal motor axons could often degenerate from the neuromuscular junctions (NMJs) in ALS. Therefore, the ability for motor axons to regrow or reconnect to NMJs is also a marker of early-stage ALS phenotype. To determine if loss-of-function mutations of TBK1 impaired motor axonal repair after injury, we seeded hMNs in microfluidic devices that allowed motor axons to sprout from the neuronal cell body chamber to a separate axonal chamber (Figure 6E). Motor neurons were plated for seven days in the soma side, and axons sprouted into the axon side through the microchannels of the device (Figure 6D-F). Motor axons were severed after seven days of culture on the device, and axonal regeneration was measured 24 hr after the axotomy (Figure 6D and 6F). TBK1^−/−^ motor neurons showed significantly reduced axonal length after injury (p<0.0001) (Figure 6G). Notably, Lys05 treatment also caused reduced axonal regeneration after injury in control hMNs (Figure S5A-C), suggesting that the reduced axonal repair might be a phenotypic consequence of lysosomal dysfunction induced TDP-43 pathology. Our results indicated that loss of TBK1 activity caused hyperexcitability and deficient axonal repair in hMNs, two key disease phenotypes found in ALS.

### Restoring TBK1 expression rescues TDP-43 pathology and neurodegeneration

Our results thus far suggest that eliminating TBK1 expression in hMNs disrupted TDP-43 homeostasis, endo-lysosomal pathway, and caused impaired homeostatic functions of hMNs. To reassure these phenotypes were the functional results of TBK1 ablation, we restored TBK1 expression by introducing the inducible TRE3G vector system containing the full-length TBK1 cDNA into the TBK1^−/−^ cells (Kim et al., 2016) (Figure 7A). Upon doxycycline (dox) induction, we found that TBK1 and p-TBK1 expression was partially restored (about 50%) with P62 levels reduced in TBK1^−/−^ motor neurons by protein blot analysis (Figure 7B, 7C, and S7A). To determine whether TBK1 restoration could restore lysosomal function, we performed flow cytometry to quantify lysosomal acidification levels using LysoTracker Red (Figure 7D). TBK1 restoration completely rescued the acidification level of lysosomes (Figure 7E), highlighting the critical role of TBK1 in maintaining lysosomal functions in motor neurons. In addition, we examined the number of RAB5-positive vesicles, which we found to be accumulated in TBK1^−/−^ motor neurons, upon TBK1 restoration (Figure 7F). Restoration of TBK1 decreased RAB5 vesicle numbers and increased lysosomal enzymatic activity (Fig. 7G, S7F, and S7G), suggesting the impaired endosomal maturation and endo-lysosomal pathway have been rescued by TBK1 restoration.

**Figure 7.**
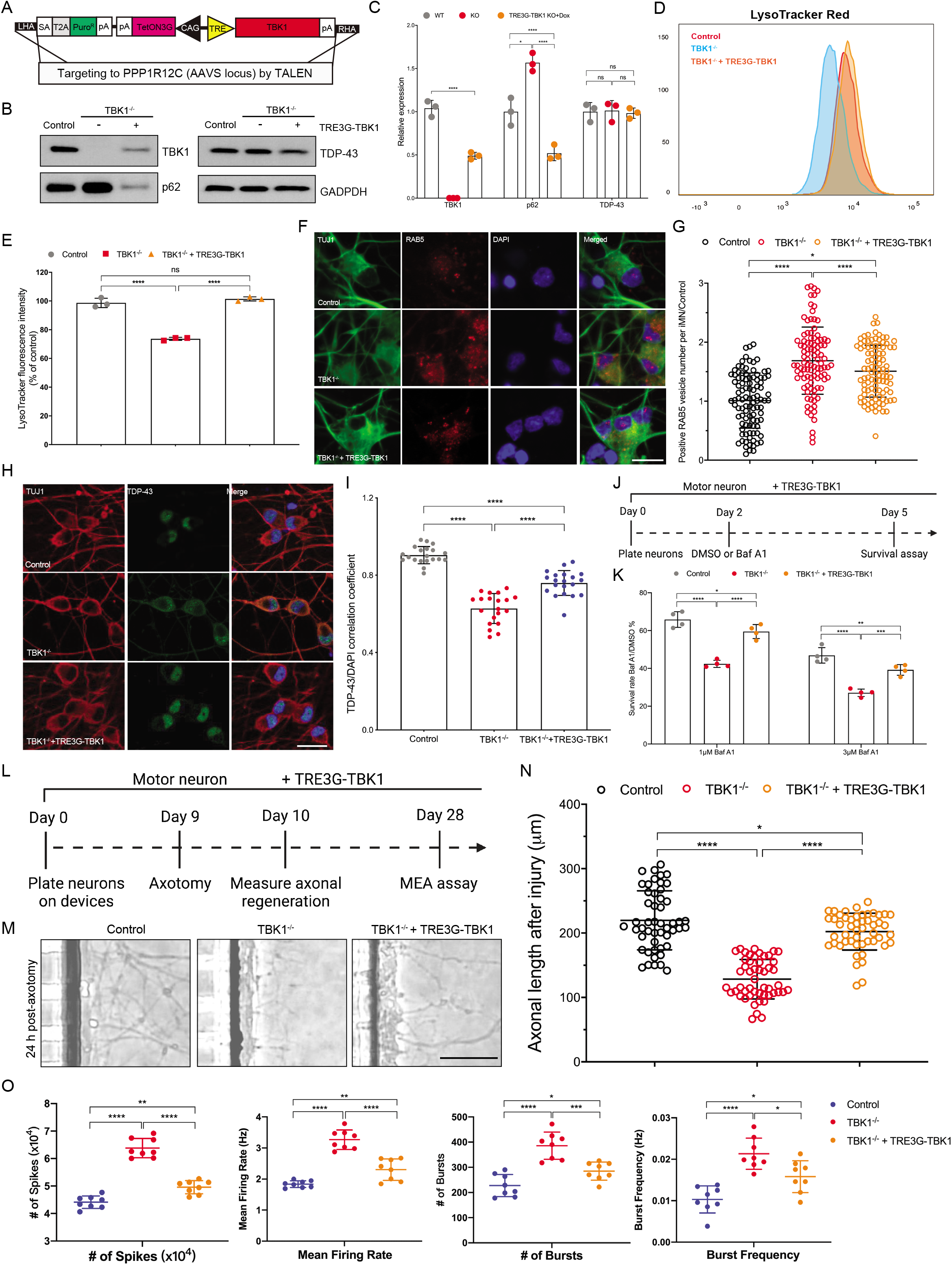
TBK1 protein restoration in TBK1^−/−^ hMNs rescues lysosomal function and attenuates motor neuron degeneration phenotypes. (A) Strategy used to insert full-length WT-TBK1 gene at the AAVs locus by TALEN. The TRE-TetON3G (TRE3G) system is used for doxycycline-inducible TBK1 expression. (B) Western blot analysis of TBK1, p62, and TDP-43 expression in control, TBK1^−/−^, and TBK1^−/−^ + TRE3G-TBK1 hMNs. (C) Quantifications of TBK1, p62, and TDP-43 levels from (B) are shown. GAPDH is used as the loading control for normalization. (D) Flow cytometry analysis of LysoTracker Red staining in control, TBK1^−/−^, and TBK1^−/−^ + TRE3G-TBK1 hMNs. (E) Quantifications of the intensity of LysoTracker Red in control, TBK1^−/−^, and TBK1^−/−^ + TRE3G-TBK1 hMNs from (D) are shown. (F) Representative images of control, TBK1^−/−^, and TBK1^−/−^ + TRE3G-TBK1 hMNs for RAB5^+^ vesicles. All cells are stained by DAPI and neuronal marker TUJ1. Scale bar, 20 μm. (G) Quantifications of the positive vesicle numbers in TBK1^−/−^ and TBK1^−/−^ + TRE3G-TBK1 hMNs from (F) are shown. Quantifications are normalized to control hMNs. (H) Representative images of control, TBK1^−/−^, and TBK1^−/−^ + TRE3G-TBK1 motor neurons for quantification of TDP-43 localization. All cells are stained by DAPI, TDP-43, and neuronal marker TUJ1. Scale bar, 20 μm. (I) Quantifications of the TDP-43/DAPI correlation coefficient from (H) are shown. (J) Experimental outline of cell survival assays in hMNs. Motor neurons are differentiated, and DMSO or 100 mM Baf A1 is added on Day 2. Cell survival assay is performed on Day 5. (K) The number of living cells is measured on Day 5 for survival assay. The surviving rate of cells treated with Baf A1 is normalized to the DMSO control. (L) Experimental outline of axotomy and MEA assays in hMNs. (M) Representative images of axonal sprouting in control, TBK1^−/−^, and TBK1^−/−^ + TRE3G-TBK1 hMNs from both pre- and post-axotomy conditions. Scale bar: 500 μm. (N) Quantifications of axonal regrowth length for control, TBK1^−/−^, and TBK1^−/−^ + TRE3G-TBK1 hMNs are shown. (O) Analysis of MEA recordings on Day 28 in culture. Compared to TBK1^−/−^ motor neurons, spike numbers and mean firing rate are reduced in TBK1^−/−^ + TRE3G-TBK1 motor neurons. Data represent means ± SE (n ≥ 3; ns means p ≥ 0.05; *p < 0.05; **p < 0.01; ***p < 0.001; ****p < 0.0001; ANOVA).

Next, we examined whether restoring TBK1 could alleviate TDP-43 pathology and early-stage neurodegenerative phenotypes in TBK1^−/−^ motor neurons. We performed ICC analysis using endogenous TDP-43 antibody and calculated the correlation coefficient of TDP-43 to DAPI staining in control, TBK1^−/−^, and the TBK1 restoration (TBK1^−/−^ + TRE3G-TBK1) motor neurons (Figure 7H). Despite the loss of TBK1 activity in the mutant hMNs, subsequent restoration of TBK1 expression attenuated the cytoplasmic localization of TDP-43 (p<0.0001) (Figure 7I). Subcellular fractionation analysis also confirmed that the pellet fraction of TDP-43 was reduced with TBK1 restoration (p<0.0001), and the nuclear fraction of TDP-43 was increased (p<0.0001) (Figure S7B and S7C). In addition, restoring TBK1 activity completely rescued the insoluble TDP-43 accumulations in TBK1^−/−^ hMNs (Figure S7D and S7E). Our results show that the expression level of TBK1 negatively correlates with the TDP-43 pathology, and TDP-43 pathology is a functional consequence of TBK1 deficiency.

We next tested whether decreased motor neuron survival upon Baf A1 treatment and the early-stage neurodegenerative phenotypes found in TBK1^−/−^ motor neurons could be alleviated by restoring TBK1. To do this, we treated motor neurons with DMSO or 100 nM Baf A1 on Day 2, and cell survival assay was performed on Day 5 (Figure 7J). Remarkably, although Baf A1 treatment sensitized TBK1^−/−^ motor neurons to neuronal death, partial restoration of TBK1 significantly rescued the motor neuron survival (p<0.0001) (Figure 7K). Thus, restoration of functional TBK1 activity protected the vulnerable motor neurons from degeneration under stress. In addition, axotomy experiment was conducted on Day 9 after motor neuron differentiation and MEA recordings were tested on Day 28 (Figure 7L). Restoration of TBK1 promoted motor neuron’s ability to regenerate after axotomy in TBK1^−/−^ motor neurons (p<0.0001) (Figure 7M and 7N). In addition, the mean firing (p<0.0001) and bursting frequency (p<0.05) were reduced in the TBK1^−/−^ hyperexcitable motor neurons (Figure 7O). Therefore, restoration of TBK1 rescues endo-lysosomal functions, attenuates TDP-43 pathology, and protects motor neuron survival in response to stress.

### Lysosomal stress sensitizes TBK1 patient-derived hMNs to neurodegeneration

Our results suggested the crucial role of TBK1 in maintaining TDP-43 homeostasis and motor neuron health and showed that complete loss of TBK1 is sufficient to drive ALS-associated phenotypes in hMNs. However, homozygous mutations of *TBK1* are embryonic lethal, and *TBK1* is haploinsufficient in ALS and FTD patients (Freischmidt et al., 2015). To determine whether haploinsufficiency of TBK1 in patient hMNs was sufficient to recapitulate ALS disease process, we generated induced pluripotent stem cells (iPSCs) from two ALS patients (MGH-138 and MGH-142) with *TBK1* mutations and induced them into motor neurons (Figure 8A). Sanger sequencing confirmed the missense mutation of p.Leu306Ile (L306I) and deletion of p.Glu463Serfs (E463Sfs) in MGH-138a and MGH-142a, respectively (Figure S8A). Consistent with a previous report (Ye et al., 2019), mutation of 138a, which was located at the kinase domain (KD) and might result in reduced kinase activity, did not show reduction in TBK1 protein level compared to healthy controls (n=4, control iPSCs: MGH-11a, 15b, 17a, and 18a) (Figure 8B and 8C). Mutation of 142a on the CCD1 domain, which was associated with protein instability, led to about 50% reduction of TBK1 proteins compared to controls (Figure 8B and 8C). Both 138a and 142a hMNs showed increased P62 levels, indicating impaired autophagy flux in both TBK1 mutant lines (Figure 8B and 8C). Therefore, our data confirmed the heterozygous mutations of *TBK1* at different locus in both cell lines, and reduced TBK1 activity was associated with aberrant autophagy activity.

**Figure 8.**
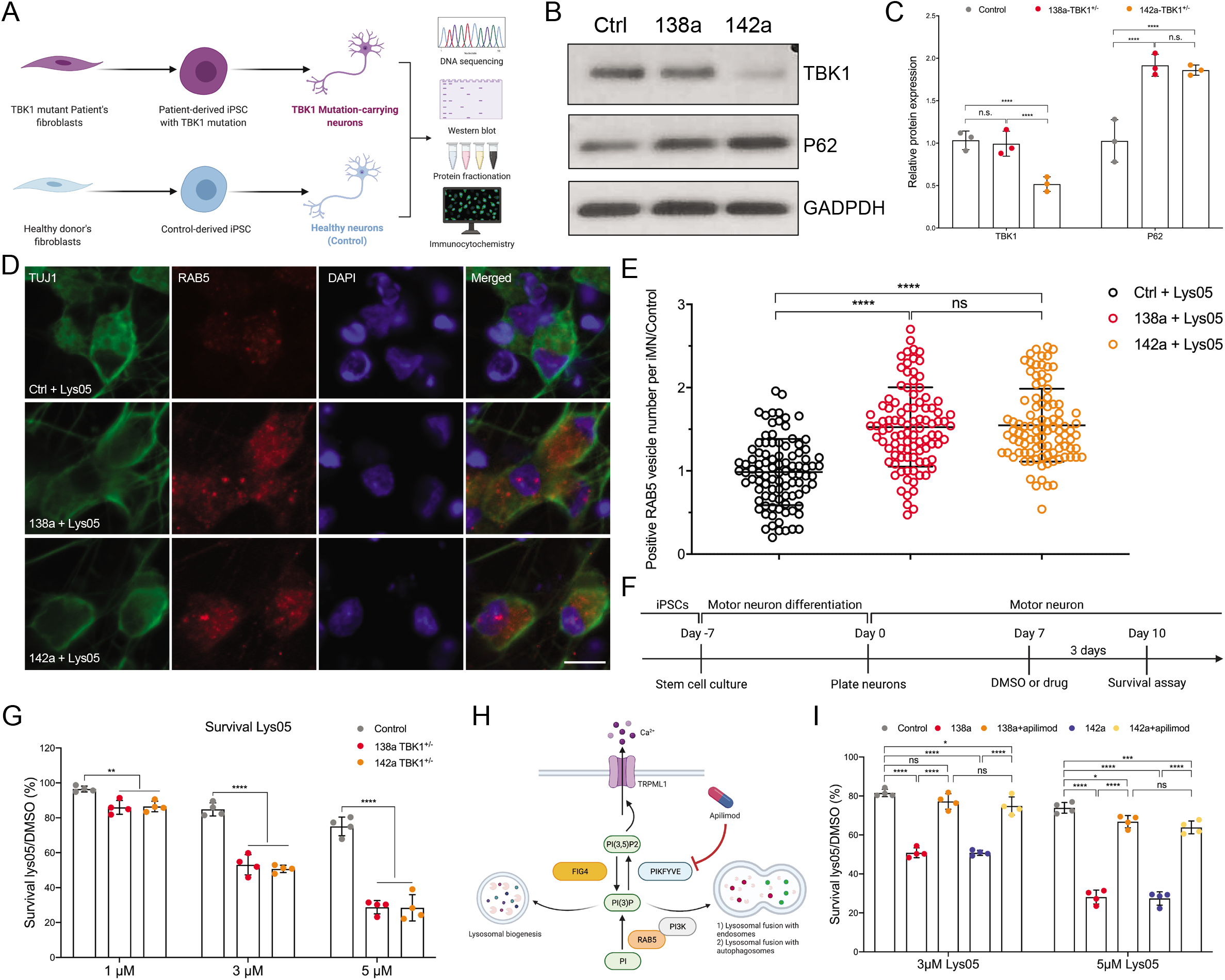
Small molecule modulator of lysosomal functions rescues the neurodegeneration of TBK1 patient hMNs under stress. (A) Schematic illustration showing the generation of iPSCs and induced hMNs from TBK1 mutant patients and healthy controls. (B) Western blot analysis of TBK1 and p62 expression in healthy controls and TBK1 patient (138a and 142a) hMNs. (C) Quantifications of TBK1 and p62 expression levels from (B) are shown. GAPDH is used as the loading control for normalization. (D) Representative images of control, 138a, and 142a hMNs treated with Lys05 for vesicle number quantification of RAB5. All cells are stained by DAPI and neuronal marker TUJ1. Scale bar, 20 μm. (E) Quantifications of the positive RAB5 vesicle numbers in TBK1^−/−^ hMNs from (D) are shown. (F) Experimental outline of cell survival assay for control and TBK1 mutant hMNs. Motor neurons are plated for 7 days and treated with Lys05 or DMSO for 72 hours. The number of living cells is measured for survival assay. (G) The control and TBK1 patient hMNs are treated with different concentrations of Lys05, and percentage of survival rate is compared to DMSO control. (H) The mechanistic activity of Apilimod in regulating vesicle trafficking and lysosomal functions. (I) The survival effect of Apilimod on TBK1 patient hMNs under lysosomal stress. The percentage of survival rate is compared to DMSO control. Data represent means ± SE (n ≥ 3; ns means p ≥ 0.05; *p < 0.05; **p < 0.01; ***p < 0.001; ****p < 0.0001; ANOVA).

To determine if reduced TBK1 activity recapitulate motor neuron degeneration in ALS processes, we examined the survival of patient and control hMNs. Consistent with our previous results, haploinsufficiency of TBK1 in patient-derived hMNs did not affect neuronal survival in culture (Figure S8B). In addition, TDP-43 mostly localized in the nuclear fraction in TBK1 patient-derived hMNs (Figure S8C), suggesting that TBK1 haploinsufficiency did not trigger TDP-43 pathology. Given that loss of TBK1 activity triggers TDP-43 and motor neuron pathology by lysosomal deficiency, we stimulated the motor neuron cultures with lysosomal stress (1 μM Lys05, 48 hr treatment). This treatment initiated a significant increase of RAB5-positive vesicles in patient-derived, but not in control, hMNs (p<0.0001) (Figure 8D and 8E), suggesting that the reduced TBK1 activity in patient-derived hMNs led to deficient endosomal maturation under lysosomal stress. In addition, we examined the motor neuron survival under lysosomal stress (Figure 8F). Strikingly, while Lys05 treatment had little effect on the survival of control hMNs, it initiated a robust degenerative response in TBK1 patient-derived hMNs in a dose-dependent manner (1 μM Lys05: p<0.01, 3 μM Lys05: p<0.0001, 5 μM Lys05: p<0.0001) (Figure 8G). To confirm the link between lysosomal functions and patient motor neuron degeneration, we use a small molecule, Apilimod, for targeting endosomal maturation and lysosomal functions. Apilimod is a PIKFYVE kinase inhibitor, which inhibits the conversion of phosphatidylinositol 3-phosphate (PI3P) into phosphatidylinositol (3,5)-bisphosphate (PI(3,5)P2) (Cai et al., 2013). PI3P anchors EEA1 to early endosomes to promote endosomal maturation and drives the fusion of lysosomes with autophagosomes and endosomes (Ikonomov et al., 2006; Lemmon, 2008). Thus, the inhibition of PIKFYVE kinase could promote endo-lysosomal functions and may compensate for reduced TBK1 activity-mediated motor neuron pathology. To determine whether PIKFYVE inhibition could rescue TBK1 patient motor neuron survival by restoring endosomal maturation and lysosomal function under stress, we examine the neuronal survival with and without Apilimod treatment. Apilimod almost completely rescued patient-derived motor neuron survival during Lys05 treatment (138a + Apilimod of 3 μM Lys05: p=0.3301, 142a + Apilimod of 3 μM Lys05: p=0.0453, 138a + Apilimod of 5 μM Lys05: p=0.0295, 142a + Apilimod of 5 μM Lys05: p=0.001) (Figure 8I). In addition, electrophysiological activity was reduced by Apilimod treatment in patient-derived motor neurons under lysosomal stress (mean firing rate: p<0.01, burst frequency: p<0.001) (Figure S8D). Therefore, small molecule inhibition of PIKFYVE kinase rescues ALS-associated disease processes induced by TBK1 haploinsufficiency.

## Discussion

The advance of next-generation sequencing techniques has led to the discovery of multiple new ALS genes (Cirulli et al., 2015), and the heterozygous loss-of-function mutations of *TBK1* have been reported to associate with ALS and FTD (Freischmidt et al., 2015). However, evaluation of the loss-of-function mutations of *TBK1* is halted by the incomplete understanding of the cell-autonomous functions of TBK1 related to motor neuron pathology (Freischmidt et al., 2017). Here we first generated a human stem cell model with homozygous loss-of-function mutations of *TBK1*. Our results indicate that loss of TBK1 activity leads to autophagy dysfunction, a key mechanism for disrupted protein degradation and ALS pathology (Beckers et al., 2021; Ramesh and Pandey, 2017). Notably, our results show that increased cytoplasmic TDP-43 protein and its insoluble accumulations, a universal pathological hallmark in ALS, is found in TBK1^−/−^ motor neurons but not in hESCs, suggesting that TDP-43 pathology may be a motor neuron- or neuron-specific phenotype, at least in the scenario of TBK1 deficiency.

TDP-43 pathology is a universal hallmark and a prominent feature of ALS. Previous studies have led to the proposal that cytosolic and insoluble accumulations of TDP-43 were initiated by autophagy or proteasome activity inhibition (Bose et al., 2011; Scotter et al., 2014). Our data suggests that proteasome inhibition reduces the soluble fraction of TDP-43 but has little effect on its insoluble accumulations. In addition, autophagy inhibition by ATG7 deletion does not cause insoluble TDP-43 accumulations or TDP-43 mislocalization, suggesting pathways independent of autophagy and proteasome activity in TBK1^−/−^ induced TDP-43 pathology. Interestingly, autophagy inhibition exacerbated the insoluble TDP-43 accumulation and reduced motor neuron survival in TBK1^−/−^ motor neurons, indicating that autophagy has a supplemental but not a dominant role in the clearance of pathological TDP-43 induced by loss of TBK1. Therefore, in contrast to the previous recognition that autophagy inhibition is a disease-causing mechanism of TBK1 mutations (Catanese et al., 2019; Oakes et al., 2017), our data provides the first direct evidence that disrupted autophagy can only exacerbate but not initiate TDP-43 pathology in hMNs.

To the best of our knowledge, our data shows the differential phospho-proteome expression map of TBK1^−/−^ motor neurons for the first time. The phospho-proteome data establishes a mechanistic link between TDP-43 pathology and the impaired endosomal pathways. Our results show impaired endosomal maturation with a reduced number of late endosomes and accumulated early endosomes in TBK1^−/−^ motor neurons, possibly through insufficient phosphorylation of RAB GTPase controlling this process. A recent study has linked increased TDP-43 aggregation with impaired endocytosis, while enhanced endocytosis reversed TDP-43 toxicity and motor neuron dysfunctions (Liu et al., 2017). In an independent study, reduced TDP-43 expression decreased the number and motility of recycling endosomes, specifically in human iPSC-derived neurons, while overexpression of TDP-43 showed opposite effects (Schwenk et al., 2016). Consistent with previous results (Leibiger et al., 2018), we show that endosomal pathways dominate the clearance of pathological TDP-43, while autophagy contributes to the clearance of TDP-43 only when the endosomal pathway is dysfunctional. In addition, the observation that loss-of-function mutations of other ALS genes also encode components of endosomal pathways, e.g., CHMP2B, highlighted the importance of the endosomal pathway in TDP-43 pathology (Leibiger et al., 2018).

Impaired endosomal maturation could lead to deficient lysosomal functions. Our results indicate reduced acidification of lysosomes, which could be a functional consequence of impaired endosome and/or autophagosome maturation, in TBK1^−/−^ motor neurons. The formation of lysosomes is accomplished by the fusion of transport vesicles from the *trans* Golgi network with endosomes and/or autophagosomes (Hu et al., 2015). During endosomal maturation, which plays a critical role in delivering lysosomal acid hydrolases to the lysosomal lumen, the internal pH can be lowered to about 4.5 in neurons (Ishida et al., 2013). The fusion of late endosomes with lysosomes activates hydrolase precursors, and mannose-6-phosphate residues, which are recognized by mannose-6-phosphate receptors, targets acid hydrolases to lysosomes (Pillay et al., 2002). Therefore, impaired endosomal fusion could result in insufficient lysosomal acidifications, which would cause subsequent lysosomal dysfunction.

Notably, our data bridges the link between TDP-43 pathology and lysosomal dysfunction. Inhibition of lysosomal activity is sufficient to cause cytosolic and insoluble TDP-43 accumulations in hMNs. Interestingly, the reduced expression of *C9ORF72* in hMNs has also been reported to show perturbation of vesicle trafficking and reduced lysosomal biogenesis, resulting in lysosomal dysfunctions (Shi et al., 2018). Previous studies also implicate several ALS or FTD mutations linked to lysosomal dysfunction, e.g., *C9ORF72*, TARDBP, CHMP2B, OPTN, VCP, and SQSTM1/p62 (Root et al., 2021). Although mechanistically divergent, our findings highlight a phenotypic convergence of TBK1 with a large portion of other genetic causes of ALS or FTD.

Consistent with previous animal experiments (Brenner et al., 2019; Gerbino et al., 2020), we find that loss of TBK1 in hMNs does not display overt motor neuron death. However, our results reveal some early-stage ALS symptoms in TBK1^−/−^ hMNs. Specifically, loss of TBK1 activity leads to motor neuron hyperexcitability, an early pathogenic mechanism occurring before molecular or anatomical symptoms of neurodegeneration in ALS. The motor neuron hyperexcitability found in TBK1 mutant hMNs could be explained by at least two mechanisms. First, reduced TBK1 activity causes impaired autophagy, which has been found to associate with neuronal hyperexcitability (McMahon et al., 2012). Similarly, in Atg7-deficient neurons, lysosomal delivery of endocytosed Kir2 channels is disrupted, and the Kir2 channels are elevated on the plasma membrane with reduced activity (Lieberman et al., 2020). These inactivated channels result in decreased Kir2 currents in the absence of autophagy, correlating with the intrinsic hyperexcitability caused by autophagy deficiency (Lieberman et al., 2020). Second, our results suggest that loss of TBK1 results in TDP-43 pathology, which is associated with deficits in RNA processing. Previous studies have revealed that mislocalization or dysfunction of TDP-43 drives intrinsic hyperexcitability of both cortical and motor neurons (Devlin et al., 2015; Dyer et al., 2021). Although the detailed mechanisms of TDP-43 dysfunction induced hyperexcitability remained unknown, possible interpretations include perturbations in ion channels and reduction in the magnitude of voltage-activated currents caused by deficits in RNA processing. In addition, we have previously identified the impaired regeneration of motor axons caused by siTDP-43 (Klim et al., 2019). Loss of TBK1 also induces the impaired axonal repair upon axotomy, a marker of early-stage ALS phenotype. Thus, we establish a link between TBK1 activity and TDP-43 pathology. Although the loss of TBK1 does not cause motor neuron death, the induced TDP-43 pathology could be linked to other early-stage disease symptoms, like hyperexcitability and impaired axonal repair in hMNs.

Our results highlight the importance of TBK1 activity in TDP-43 and motor neuron pathology, making TBK1 a key therapeutic target for ALS and FTD patients with reduced TBK1 activity. We establish a new approach for rescuing the TDP-43 pathology and neurodegeneration induced by reduced TBK1 function: restoring or replacing TBK1 activity. Our approach restores only partial expression of TBK1, but we have observed significantly reduced TDP-43 pathology and alleviated neurodegenerative phenotypes. Although whether overdosed TBK1 will cause any detrimental effects on control or mutant motor neurons remains unknown and needs further validation, our approach provides information about the effects of forced TBK1 expression in mutant cells, which could support the development of therapeutic strategies for TBK1 mutant ALS patients.

Our results show that loss of TBK1 in motor neurons is related to TDP-43 pathology and neurodegenerative phenotypes. However, haploinsufficiency of TBK1 or *Tbk1* deletion in mice normally do not display any disease phenotype (Gerbino et al., 2020). We generated human patient iPSCs and induced hMNs harboring *TBK1* mutations at different locus to address the clinical relevance of reduced TBK1 activity and ALS disease processes. Our results show that TBK1 patient hMNs is more sensitive to lysosomal stress, highlighting the role of lysosomal activity in ALS processes. Previous studies have implicated decreased lysosomal activity with age (Carmona-Gutierrez et al., 2016; Sun et al., 2020), and our results may link certain neurodegenerative diseases, like TBK1 mutant associated ALS/FTD, to aging. In addition, the fact that PIKFYVE inhibitors, which could reverse *C9ORF72* neurodegeneration (Shi et al., 2018), rescued the neurodegeneration phenotype in TBK1 patient hMNs under lysosomal stress may provide novel insights into the therapeutics of ALS and FTD or other neurodegenerative disorders.

## Supporting information

Graphic abstract

## Author contributions

J.H. and K.E. initiated and designed the project. J.H. performed most of the experiments. J.H., M.F.W., and K.E. wrote the manuscript and interpreted the data. G.N. helped the cell culture and western blot experiments. I.G.J. and F.L. helped the microfluidic device, axotomy experiments, and imaging analysis. A.F. helped molecular cloning and generating the TBK1 inducible system. M.L. participated the flow cytometry experiments. B.J. helped MEA experiments. M.Q. and D.A. participated in some of the cell survival assay and imaging experiments. B.B. performed phosphoproteomics and analyzed the data. Z.D. provided shATG7 and Tet-pLKO-puro plasmid constructs. K.E. supervised this project.

## Acknowledgements

We want to thank Dr. Lee Rubin, Dr. Clifford Woolf, and members of the Eggan lab for their insightful discussions and advice on this work. We thank Amie Holmes, Susi Jakob and the SCRB15 class at Harvard University for providing support for this project. We thank the Harvard Center for Biological Imaging for infrastructure and support. Support to K.E. was provided by Target ALS, NIH 5R01NS089742 and the Harvard Stem Cell Institute.

## Supplementary figures

**Figure S1. Validation of TBK1 deletion in hESCs, related to Figure 1.** (A) Sanger sequencing of control and two candidate mutant clones. (B) TIDE analysis of indels percentage of the two candidate clones based on the sequencing data. Deletions were shown on the left X axis, and insertions were shown on the right based on the number of deleted or inserted bps. (C) Immunostaining of TBK1 for hESC cultures of control and TBK1^−/−^ cells. Nucleus were stained for DAPI. Scale bar, 20 μm. (D) Subcellular fractionation analysis of Lamin B1 (pellet control) and TDP-43 levels in different fractions of control and TBK1^−/−^ hESCs. Representative blots show TDP-43 levels in cytosol extract (CE), membrane extract (ME), nuclear extract (NE), and pellet extract (PE).

**Figure S2. Induction of motor neuron using NGN2 programming, related to Figure 2.** (A) Design of lentiviral vectors for NGN2-induced motor neuron differentiation through Dox-inducible promotor (TetO) mediated expression. Cells were transduced with the two viruses and selected with puromycin. The induction of motor neuron differentiation was based on NGN2 expression and motor neuron patterning. (B) Representative images showing the Hb9::GFP positive cells in control and TBK1^−/−^ cells on Day 7. (C) Immunoblot analysis of TBK1, P62, and TDP-43 protein expression for two candidate mutant clones. Quantifications of the protein levels from 3 independent experiments are shown. GAPDH was used as an internal control. (D) Immunostaining of motor neuron marker, ISL-1, in control and TBK1^−/−^ cells. All cells are stained by DAPI and neuronal marker TUJ1. Scale bar, 20 μm. (E) Quantifications of percentage of cells expressing ISL-1 are shown.

**Figure S3. Proteasome inhibition by MG132 does not increase insoluble TDP-43 accumulation or motor neuron death, related to Figure 3.** (A) Western blot analysis of control and TBK1^−/−^ hMNs treated with DMSO or MG132. Accumulations of TDP-43 proteins are shown in the soluble (RIPA) and insoluble (urea) fractions. Quantification of TDP-43 expression between DMSO and MG132 treated hMNs was shown. GAPDH in the soluble fraction is used as the loading control for normalization. (B) Experimental outline of motor neuron survival assay. Motor neurons were differentiated and plated and were treated with DMSO or drug on Day 2 for 48 hr before the assay. (C) The survival rate of control and TBK1 ^−/−^ hMNs in response to different concentration of MG132 and Baf A1. The relative survival rate was compared to that of DMSO control. (D) Effects of shATG7 on LC3-II levels and degradation during autophagic maturation. The shNTC control and shATG7 hMNs are treated or not treated with BafA1 to inhibit autophagic degradation of LC3-II. (E) Immunostaining of ISL-1 in shNTC and shATG7 hMNs. All cells are stained by DAPI and neuronal marker TUJ1. Scale bar, 20 μm. (F) Quantifications of percentage of cells expressing ISL-1 are shown.

**Figure S4. Loss of TBK1 in hMNs impairs endosomal maturation and lysosomal function, related to Figure 4.** (A) Schematic illustration showing the potential function of TBK1 in endosomal maturation. TBK1 promotes the maturation of early endosomes into late endosomes by converting RAB5 to RAB7. (B) Representative images of control and TBK1^−/−^ motor neurons for vesicle number quantification of LAMP1. All cells are stained by DAPI and neuronal marker MAP2. Scale bar, 20 μm. (C) Quantifications of the positive LAMP1 vesicle numbers in TBK1^−/−^ hMNs from (B) are shown. All quantifications are normalized to control hMNs.

**Figure S5. Lysosomal inhibition leads to impaired axonal regeneration in control hMNs, related to Figure 5.** (A) Schematic of axonal regrowth experiment upon injury. The control hMNs are plated on the left chamber (soma side) of the device, and axons sprout into the right chamber (axon side) through the channels connecting both sides. Lys05 was added 2 days prior to axotomy. Axotomy is performed by washing out the axon side to cut the sprouted axons. The ability of axonal repair is measured by the length of re-sprouted axons. (B) Representative images of axonal sprouting in control hMNs with and without Lys05 treatment from pre- and post-axotomy conditions. Scale bar: 500 μm. (C) Quantifications of axonal regrowth length for control hMNs with and without Lys05 treatment are shown. The data of individual neurites is displayed as a single dot.

**Figure S6. Loss of TBK1 does not cause motor neuron degeneration under basal conditions, related to Figure 6.** (A) Experimental outline of motor neuron survival assay in hMNs. Motor neurons were differentiated and plated on Day 0. Cell survival assay was performed on Day 2, Day 14, and Day 28. (B) Immunostaining of TUJ1 and DAPI in control and TBK1^−/−^ hMNs on Day 2 before the survival assay. (C) The survival rate of control and TBK1^−/−^ hMNs on Day 14 and Day 28 compared to that of Day 2. (D) Number of active electrodes in control and TBK1^−/−^ hMNs cultured on MEA plates. Analysis was performed at week 3 and week 4. (E) Analysis of MEA recordings for mean firing rate in TBK1^−/−^ hMNs treated with different dose of retigabine at week 4.

**Figure S7. Restoring TBK1 activity reduced insoluble TDP-43 by rescuing endo-lysosomal function, related to Figure 7.** (A) Immunoblot analysis of TBK1 and pTBK1 expression in control and TBK1−/− hMNs with and without TBK1 restoration. (B) Subcellular fractionation analysis of TBK1 (cytosol control), Lamin B1 (nuclear and pellet control), STING (membrane control), and TDP-43 levels in different fractions of control and TBK1^−/−^ hMNs with and without TBK restoration. (C) Quantifications of the partition of TDP-43 in nuclear and pellet fraction for control, TBK1^−/−^, and TBK1^−/−^ + TRE3G-TBK1 hMNs are shown. (D) Western blot analysis of control, TBK1^−/−^, and TBK1^−/−^ + TRE3G-TBK1 hMNs treated with DMSO or Lys05. Accumulations of TDP-43 proteins are shown in the soluble (RIPA) and insoluble (urea) fractions. GAPDH in the soluble fraction is used as the loading control for normalization. (E) Quantifications of insoluble TDP-43 expression between DMSO and Lys05 treated hMNs are shown. (F) Lysosome enzymatic activity assay in control, TBK1^−/−^, and TBK1^−/−^ + TRE3G-TBK1 hMNs. Self-quenched substrate is used as endocytic cargo, and fluorescence signal, generated by lysosomal degradation, is proportional to the intracellular lysosomal activity. (G) Quantifications of the relative fluorescent intensity from (F) are shown.

**Figure S8. TBK1 patient-derived hMNs does not display overt neurodegeneration, related to Figure 8.** (A) Sanger sequencing validation of MGH-138a and MGH-142a patient-derived iPSCs. (B) The survival rate of control, 138a TBK1^+/-^, and 142a TBK1^+/-^ hMNs on Day 14 and Day 28 compared to that of Day 2. (C) Subcellular fractionation analysis of Lamin B1 (pellet control) and TDP-43 levels in different fractions of control 138a TBK1^+/-^, and 142a TBK1^+/-^ hMNs. (D) Analysis of MEA recordings for mean firing rate and burst frequency in Lys05 treated patient-derived TBK1^+/-^ hMNs with and without Apilimod.

**Figure S9. Graphic abstract: Lysosomal degradation of pathological TDP-43 is mediated by TBK1.** Schematic illustration of TBK1-mediated pathological TDP-43 clearance pathway by lysosomal function. Cytoplasmic TDP-43 and their insoluble accumulations are cleared primarily by endo-lysosomal pathway in normal hMNs. Autophagy supplements to that clearance function. In hMNs with loss of TBK1 activity, impaired endosomal maturation and subsequent lysosomal dysfunction leads to the accumulation of pathological TDP-43.

## STAR Methods

### Resource Availability

#### Lead Contact

Further information and requests for resources can be directed to by the lead contact, Kevin Eggan

#### Materials availability

Cell lines generated in this study are available upon request from the lead contact, Kevin Eggan, but a completed Materials Transfer Agreement may be required.

#### Data and Code Availability

The phospho-proteomic dataset will be available on ProteomeXchange. This paper does not report original code. Any additional information required to reanalyze the data reported in this paper is available from the lead contact upon request.

## Experimental Model and Subject Details

### Human pluripotent stem cell lines

The HUES3 *Hb9::GFP* and iPSC control 11a, 15b, 17a, and 18a used in this study were previously approved by the institutional review boards of Harvard University. Patient fibroblasts of MGH-138a and 142a were obtained from Massachusetts General Hospital (MGH) and converted into iPSCs in Harvard Stem Cell Institute. Specific point mutations were confirmed by PCR amplification followed by Sanger sequencing. Our lab screens for mycoplasma contamination weekly using the MycoAlert kit (Lonza) with no cell lines used in this study testing positive. The use of these cells at Harvard was further approved and determined not to constitute Human Subjects Research by the Committee on the Use of Human Subjects in Research at Harvard University.

## Methods Details

### Cell Culture and differentiation

Pluripotent stem cells were cultured with mTeSR plus medium (Stem Cell Technologies) on Matrigel (BD Biosciences) coated tissue culture dishes. Stem cells were maintained in 5% CO_2_ incubators at 37 °C and passaged as small aggregates after 1mM EDTA treatment. After dissociation, 10 μM ROCK inhibitor (Sigma, Y-27632) was added to cell culture for 24 hr to prevent cell death, and then incubated with lentiviruses (FUW-M2rtTA, TetO-Ngn2-Puro). The HUES3 *Hb9::GFP* cell line has been previously described (Han et al., 2011). Motor neurons were differentiated using a modified protocol based on previous strategies (Amoroso et al., 2013; Nehme et al., 2018). This protocol relies on neural induction through NGN2 programming, SMAD inhibition through small molecules, and motor neuron patterning through retinoic activation and Sonic Hedgehog signaling. In brief, hESCs were dissociated into single cells using accutase (Stem Cell Technologies), and then plated on Matrigel-coated culture dish at a density of 80,000 cells per cm^2^ with mTeSR plus medium (Stem Cell Technologies) supplemented with 10 μM ROCK inhibitors (Sigma, Y-27632). When cells reached confluency, medium was changed to N2 medium (DMEM-F12 (Life Technologies) supplemented with ×1 Gibco GlutaMAX (Life Technologies), ×1 N-2 supplement (Gibco), × 0.3% Glucose) on Day 1-3. For small molecule treatment, 10 μM SB431542 (Custom Synthesis), 100 nM LDN-193189 (Custom Synthesis), 1 μM retinoic acid (Sigma), 1 μM SAG (Custom Synthesis) were added on Day 1-3. For NGN2 induction and neuron selection, 20 mg/ml Doxycycline and 10 mg/ml puromycin were added on Day 2-3. On Day 4-7, the N2 medium was changed to Neurobasal (Neurobasal (Life Technologies) supplemented with ×1 N-2 supplement (Gibco), ×1 B-27 supplement (Gibco), ×1 Gibco GlutaMAX (Life Technologies) and 100 μM non-essential amino-acids: NEAA) along with 1 μM retinoic acid, 1 μM SAG, 20 mg/ml Doxycycline, 10 mg/ml puromycin, and 10 ng/ml of neurotrophic factors: glial cell-derived neurotrophic factor (GDNF), ciliary neurotrophic factor (CNTF) and brain-derived neurotrophic factor (BDNF) (R&D). For post-mitotic cell selection, 10 mM 5-Fluoro-2’-deoxyuridine thymidylate synthase inhibitor (FUDR, Sigma) was added on Day 5-7. From Day 8, media was changed to the motor neuron maintenance media containing Neurobasal (Neurobasal (Life Technologies) supplemented with ×1 N-2 supplement (Gibco), ×1 B-27 supplement (Gibco), ×1 Gibco GlutaMAX (Life Technologies) and 100 μM non-essential amino-acids: NEAA) supplemented with 10 ng/ml of neurotrophic factors (GDNF, CNTF, and BDNF) and 10 mM FUDR. Half of the maintenance media was changed every two days.

### Cell viability assay

Motor neuron viability assay was performed based on the CellTiter-Glo luminescent for each sample according to the manufacturer’s recommendations (Promega G7570). In brief, motor neurons were plated in triplicate onto 96-well plates. After treatment incubation, motor neurons were incubated for 10 min with CellTiter-Glo reagent at room temperature, and luminescence was measured using the Cytation imaging reader (BioTek). Background luminescence was measured using medium without cells and then subtracted from experimental values.

### CRISPR/Cas9 genome editing

TBK1 mutant cell lines were generated using CRISPR/Cas9-mediated genome editing as previously described (Cong and Zhang, 2015; Klim et al., 2019). To generate loss-of-function alleles of TBK1, control hESCs were transfected with guide RNAs targeting the desired location of human TBK1 gene. TBK1 gRNAs were designed using a web tool: CHOPCHOP (https://chopchop.rc.fas.harvard.edu) (Labun et al., 2016). Guides were cloned into a vector with the human U6 promotor (custom synthesis, Broad Institute), with a BbsI cleavage site. The modified TBK1 gRNA sequences were used for Cas9 nuclease genome editing: guide 1: 5’-CACCG CAAGAATATACTAATGAGGT-3’, guide 2: 5’-CACCGATATCAAGAATATACTAATG-3’. After annealing and ligation, vectors were cloned in OneShot Top10 (ThermoFisher Scientific) cells and isolated using Qiagen MIDI-prep kit (Qiagen). After verification by sequencing, 1 μg of each vector containing the guide and 1.5 μg of the pSpCas9n(BB)-2A-Puro (PX462) V2.0 (Addgene) were added to cells for transfection using the Neon Transfection System (ThermoFisher Scientific). Colonies were picked on day 10 after transfection and genotyped by PCR amplification and sequencing. The TBK1 amplification and sequencing primers used were as followed: TBK1_Forward: 5’ GTCAGAAGAATGGATAAGGTGAG 3’, TBK1_Reverse: 5’ CAGATCTGACATCAAGTGTTACAG 3’, and TBK1_Sequencing: 5’ ATGCTTCACTAGACTGCATATC 3’. Loss-of-function mutations of TBK1 were further confirmed using immunoblot analysis.

### Immunocytochemistry and imaging

For immunocytochemistry, cells were fixed with 4% PFA for 20 min, and were subject to permeabilization with 0.25% Triton-X in PBS for 40 min. After that, cells were blocked using 10% donkey serum supplemented with 0.1% Triton-X in PBS (blocking buffer) for 1 hr at room temperature. Primary antibody diluted in blocking buffer was used for cell incubation overnight at 4 °C. At least three washes (5 min incubation each) with PBS were carried out, before incubating the cells with secondary antibodies (diluted in blocking buffer) for 1 hr at room temperature. Nuclei was stained with DAPI. The following antibodies were used in this study: TBK1 (1:200, Abcam ab109735), SQSTM1 / p62 (1:200, Abcam ab56416), TDP-43 (1:200, Proteintech Group 10782 & Cell Signaling Technology 3448S), Islet1 (1:500, Abcam ab20670), MAP2 (1:10,000, Abcam ab5392), TUJ1 (1:1,000, R&D Systems MAB NL493), EEA1 (1:100, Cell Signaling Technology 3288 & BD Biosciences 610456), RAB5 (1:100, Cell Signaling Technology 46449S), RAB7 (1:100, Cell Signaling Technology 9367S), LAMP1 (1:100, Abcam ab25630), phospho-TBK1 (1:200, Cell Signaling Technology 13498S). Secondary antibodies used (488, 555, and 647) were AlexaFluor (1:1,000, Life Technologies). Images were acquired using a Zeiss LSM 880 confocal microscope or a Nikon Eclipse Ti microscope. Images were analyzed using ImageJ or NIS--Elements (Nikon).

### Western blot assays

For protein extraction, cells were lysed using 1% SDS lysis buffer (PhosphoSolutions, 100-LYS) with a brief sonication. For insoluble protein extraction, cells were lysed using RIPA buffer (Life Technology, 89900) with protease inhibitors (Life Technology, 78425) for 20 min on ice. After centrifuge, the supernatant was collected as soluble fraction, and the insoluble fraction was collected from the pellet using 8M urea with 4% CHAPS, 40 mM Tris, and 0.2% Bio-Lyte 3/10 ampholyte (Bio-Rad, 1632103). For subcellular fractionation, cells were subject to sequential extract using the subcellular fractionation kit for cultured cell (Thermo Scientific, 78840) according to manufacturer instructions. After sample preparation, equivalent amount of 5-10 μg protein from each ample were subjected to electrophoresis using 4–20% SDS-PAGE (Bio-Rad, 4561096) and then transferred to a PVDF membrane (Thermo Fisher, 88518). The membrane was blocked with 3% Bovine Serum Albumin (Sigma, A9647) for 1 hr at room temperature, and incubated with primary antibodies overnight at 4°C. The next day, membrane was washed in TBST X3, 10 min each, and then incubated with HRP-conjugated secondary antibodies for 1 hr at room temperature. After that, membrane was washed in TBST X3, 10 min each, and then developed using ECL Western Blotting Detection System (VWR, 95038) and blotting films (Genesee Scientific, 30-810).

### TDP-43 localization assay

For analysis of correlation coefficient of TDP-43 and nuclei, motor neurons were stained for TDP-43 (Cell Signaling Technology), TUJ1 (R&D Systems), and counterstained with DAPI. TUJ1 staining was used to determine the neuronal cell body, and the Pearson’s correlation coefficient was calculated using NIS-Elements (Nikon) for TDP-43 and DAPI with at least 50 neurons being analyzed. Cells were segmented into the nuclear (DAPI positive) and cytoplasmic (DAPI negative) regions by NIS. The TDP-43 intensity in these two regions were used to determine the nuclear to cytoplasmic ratio (correlation coefficient) for TDP-43 fluorescent intensity.

### Multi-electrode array (MEA) recordings

MEA recording assay was performed according to Axion Biosystems protocols. In brief, 12-well MEA plates (Axion Biosystems, M768-GL1-30Pt200) with 64 electrodes per well were coated with PDL and laminin. Motor neurons were seeded at a density of 500,000 cells per well and were fed 2-3 times per week. Neuronal activity was measured weekly for 5 min using the Maestro 12-well 64 electrodes per well micro-electrode array (MEA) plate system (Axion Biosystems, Atlanta, GA). Spontaneous activity was recorded from all channels simultaneously using a gain of 1200x and a sampling rate of 12.5 kHz/channel. All data reflected well-wide mean from active electrodes, which were defined as having >1 spikes/min.

### Axotomy

Standard neuron microfluidic devices (SND150, XONA Microfluidics) were mounted on glass coverslips coated with 0.1 mg/ml poly-L-ornithine (Sigma-Aldrich) and 5 μg/ml laminin (Invitrogen). Motor neurons were cultured on the devices at a concentration of around 250,000 cells per device for 7 days. Axotomy was performed by repeated vacuum aspiration and reperfusion using PBS in the axonal side of the device until all axons were diminished without disturbing the soma side cells. For the experiment, two independent replicates were performed with at least 20 neurites measured.

### Lysosomal enzyme activity assay

Control and TBK1^−/−^ motor neurons were incubated with Self-Quenched Substrate (Abcam Cat No. ab234622) following manufacturer’s instructions. Briefly, cells were incubated with the substrate for 1hr and fixed with 4% PFA at room temperature for 20 min. After washing with PBS, cells were stained with DAPI and then mounted for imaging with Zeiss LSM880 Confocal Laser Scanning microscope. Images were analyzed using ImageJ. Mean fluorescence intensity was quantified in control and TBK1^−/−^ motor neurons.

### Molecular cloning and viral production

Stable cell lines expressing inducible shATG7 and GFP-mCherry-RAB5-DN were made by lentivirus infection. shATG7 hairpin sequence GGAGTCACAGCTCTTCCTTAC was from Dou lab, and was cloned into Tet-pLKO-puro ‘all-in-one’ tetracycline-inducible vector (Dou et al., 2015). pCIG3_mCherry_Rab5_DN was a gift from Felicia Goodrum (Addgene: #75111). Viruses were produced as follows. HEK293 cells were transfected with viral vectors containing genes of interest and viral packaging plasmids (pPAX2 and VSVG) with lipofectamine 3000 (Invitrogen) in Opti-MEM (Gibco). The medium was changed 24 hr, and viruses were harvested at 48 hr and 72 hr after transfection. The supernatants were filtered with 0.45 μM filters, incubated with Lenti-X concentrator (Clontech) for 24 hr, and centrifuged at 1,500 g at 4 °C for 45 min. The pellets were resuspended in 300 μl DMEM supplemented with 10% FBS and stored at −80 °C.

### Mass spectrometry

The phospho-proteomics was performed at Harvard Center for Mass Spectrometry (HCMS). Cell pellets were prepared using DF Covaris duffer and undergo protein extraction procedure for 120 s with 10% power in Covaris S220 focused ultrasound instrument (Woburn). Samples were resolubilized in 50 mM TEAB (try-ethyl ammonia bicarbonate) buffer for reduction and alkylation, further digested in buffer trypsin (Promega) for 5 hours. The digested samples were enriched by High-Select™ TiO2 Phosphopeptide Enrichment Kit (Thermo-Fisher) according to the vendor’s instructions. Enriched phosphopeptides were labeled with TMT16plexPRO (Thermo-Fisher) according to manufacturer protocol. Sample fraction was submitted for single LC-MS/MS experiment that was performed on a Lumos Tribrid (Thermo) equipped with 3000 Ultima Dual nanoHPLC pump (Thermo). The Lumos Orbitrap was operated in data-dependent mode for the mass spectrometry methods. The mass spectrometry survey scan was performed in the Orbitrap in the range of 400 –1,800 m/z at a resolution of 6 × 104, followed by the selection of the twenty most intense ions (TOP20) for CID-MS2 fragmentation in the Ion trap using a precursor isolation width window of 2 m/z, AGC setting of 10,000, and a maximum ion accumulation of 50 ms. Ions in a 10 ppm m/z window around ions selected for MS2 were excluded from further selection. The same TOP20 ions were subjected to HCD MS2 event in Orbitrap part of the equipment. The fragmentation isolation width was 0.8 m/z, AGC was 50,000, the maximum ion time was 150 ms, normalized collision energy was 38V and an activation time of 1 ms for each HCD MS2 scan was set.

### Mass spectrometry analysis

Raw data were analyzed in Proteome Discoverer 2.4 (Thermo Scientific) software with Byonic 3.5 node and ptmRS node. Assignment of MS/MS spectra was performed using the Sequest HT algorithm by searching the data against a protein sequence database including all entries from the Human Uniprot database. A MS2 spectra assignment false discovery rate (FDR) of 1% on protein level was achieved by applying the target-decoy database search. Filtering was performed using a Percolator (64bit version) (Kall et al., 2008). For quantification, a 0.02 m/z window centered on the theoretical m/z value of each the six reporter ions and the intensity of the signal closest to the theoretical m/z value was recorded. Reporter ion intensities were exported in result file of Proteome Discoverer 2.4 search engine as an excel tables. The exact place of phosphor moiety was analyzed by ptmRS program (Taus et al., 2011). Differentially expressed proteins between sample groups were analyzed using R script programs based on Bioconductor (https://www.bioconductor.org/), Statistical analysis for differentially expressed proteins was based on peptide level to find changes that are statistically significant between two sets.

### TBK1 cDNA plasmid transfection

For TBK1 rescue experiments, targeting vector with doxycycline inducible system was generated. EGFP was replaced with WT TBK1 cDNA (Addgene: #82285) in AAVS1-TRE3G-EGFP (Addgene: #52343) using MluI/SalI sites. The resulting vector was transfected together with TALEN vectors (Addgene: #52341/52342) targeting AAVS locus. TBK1^−/−^ cells were transfected using Neon Transfection System with the vector and screened by puromycin for at least 2 weeks.

### Data presentation and statistical analysis

In all figure elements, bars and lines represent the median with error bars representing standard deviation. The box and whisker plots display the minimum to maximum. Data distribution was assumed to be normal, but this was not formally tested. The statistical analyses between two groups were performed using a two-tail unpaired Student’s t-test, and differences between more than two groups were analyzed by one way--ANOVA with Tukey correction for multiple testing. Significance was assumed at P < 0.05. Error bars represent the s.d. unless otherwise stated. Analysis was performed with the statistical software package Prism 8 (Graph Pad).

