## Supplementary figures and images for "Loss of TBK1 activity leads to TDP-43 proteinopathy through lysosomal dysfunction in human motor neurons"

### Graphic abstract

#### Graphic abstract

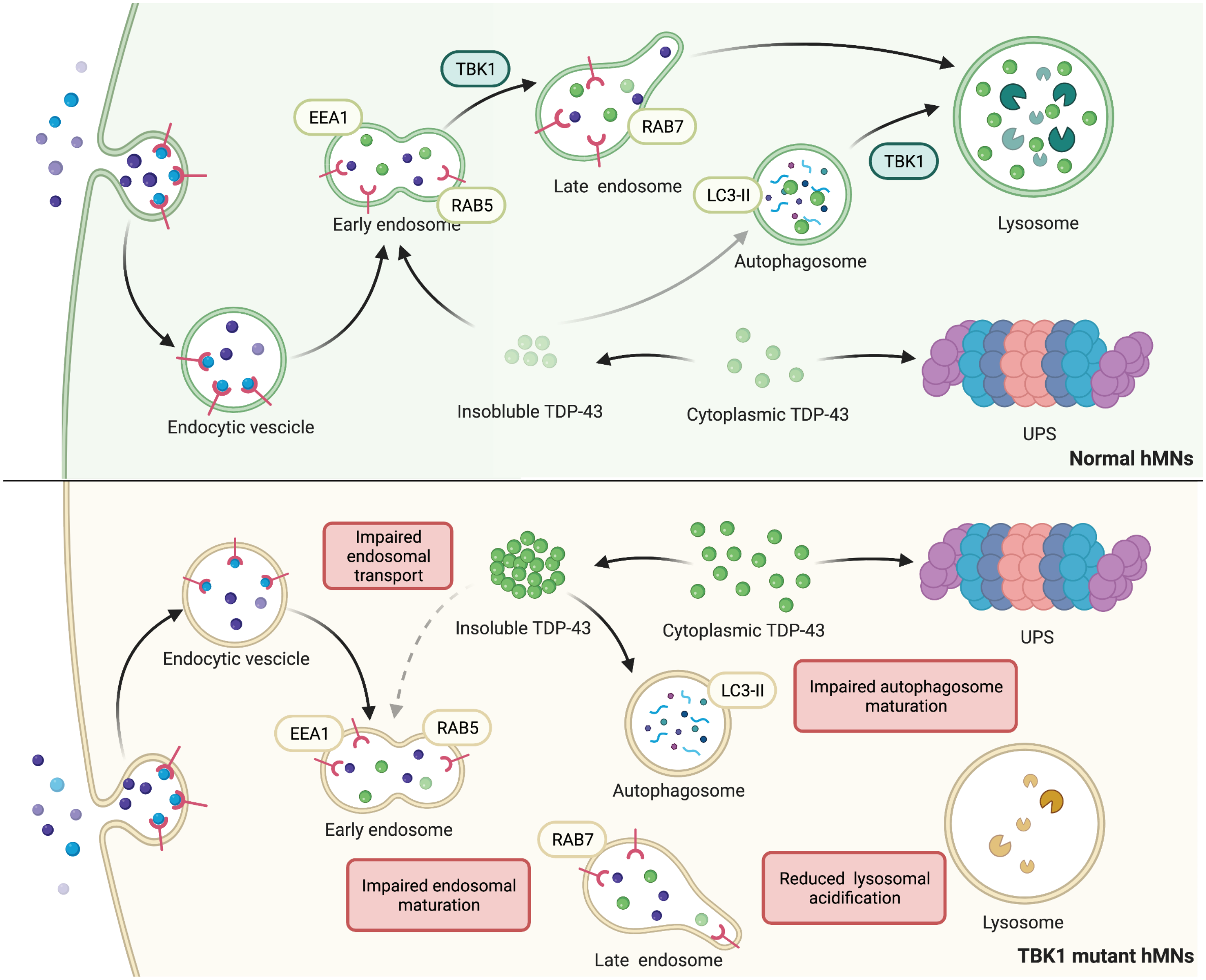

### Supplemental Figure 1

#### Supplemental Figure 1

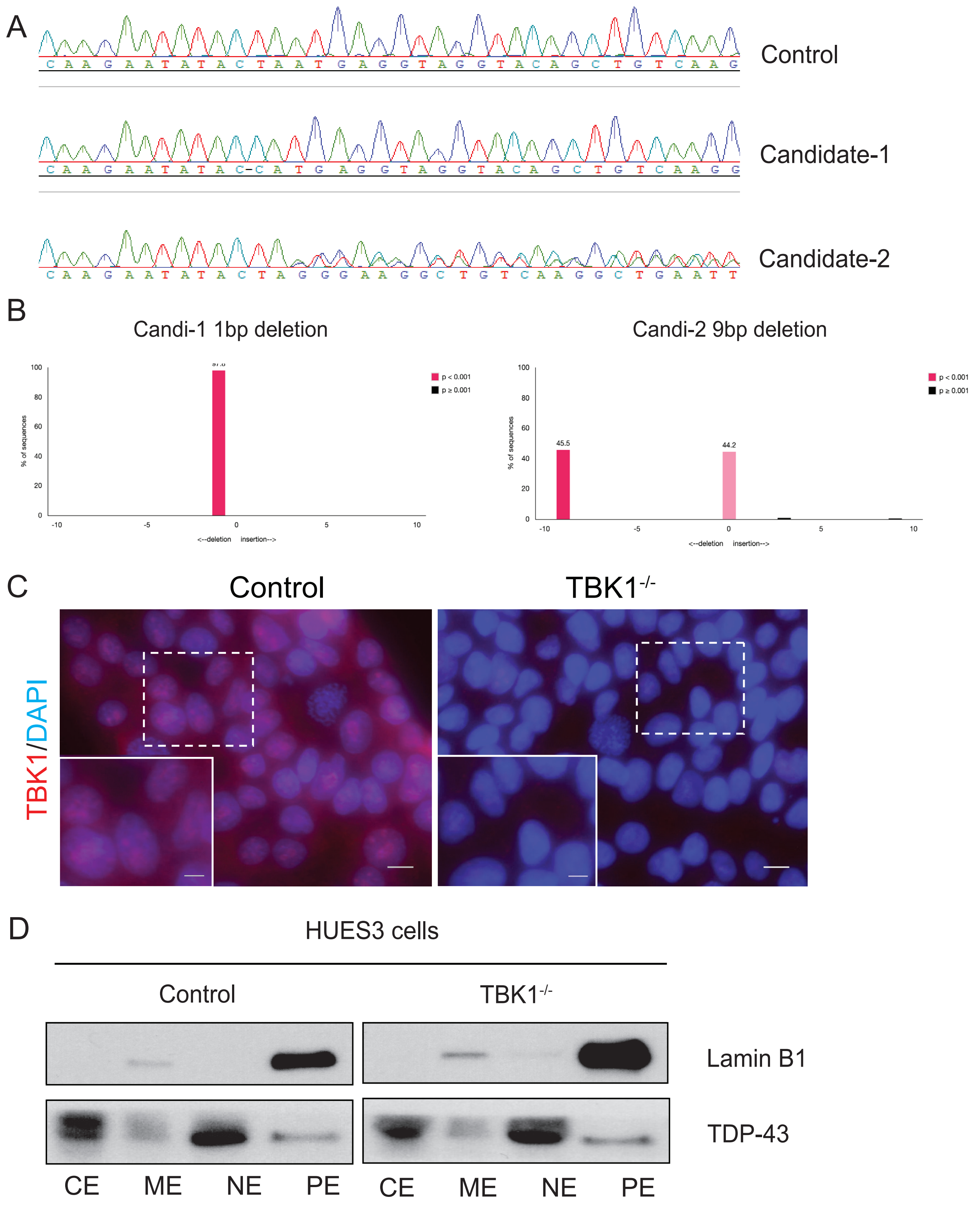

### Supplemental Figure 2

#### Supplemental Figure 2

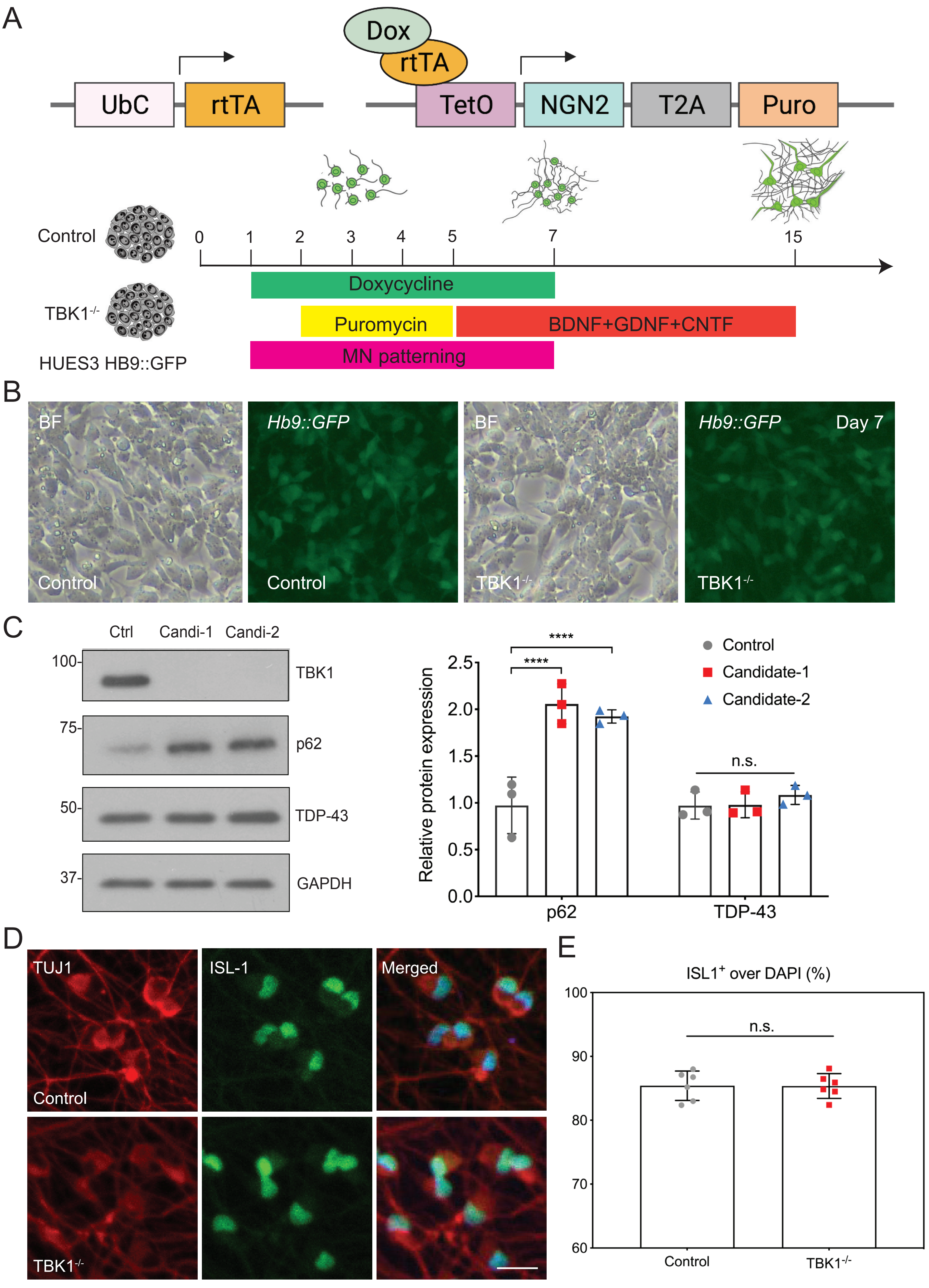

### Supplemental Figure 3

#### Supplemental Figure 3

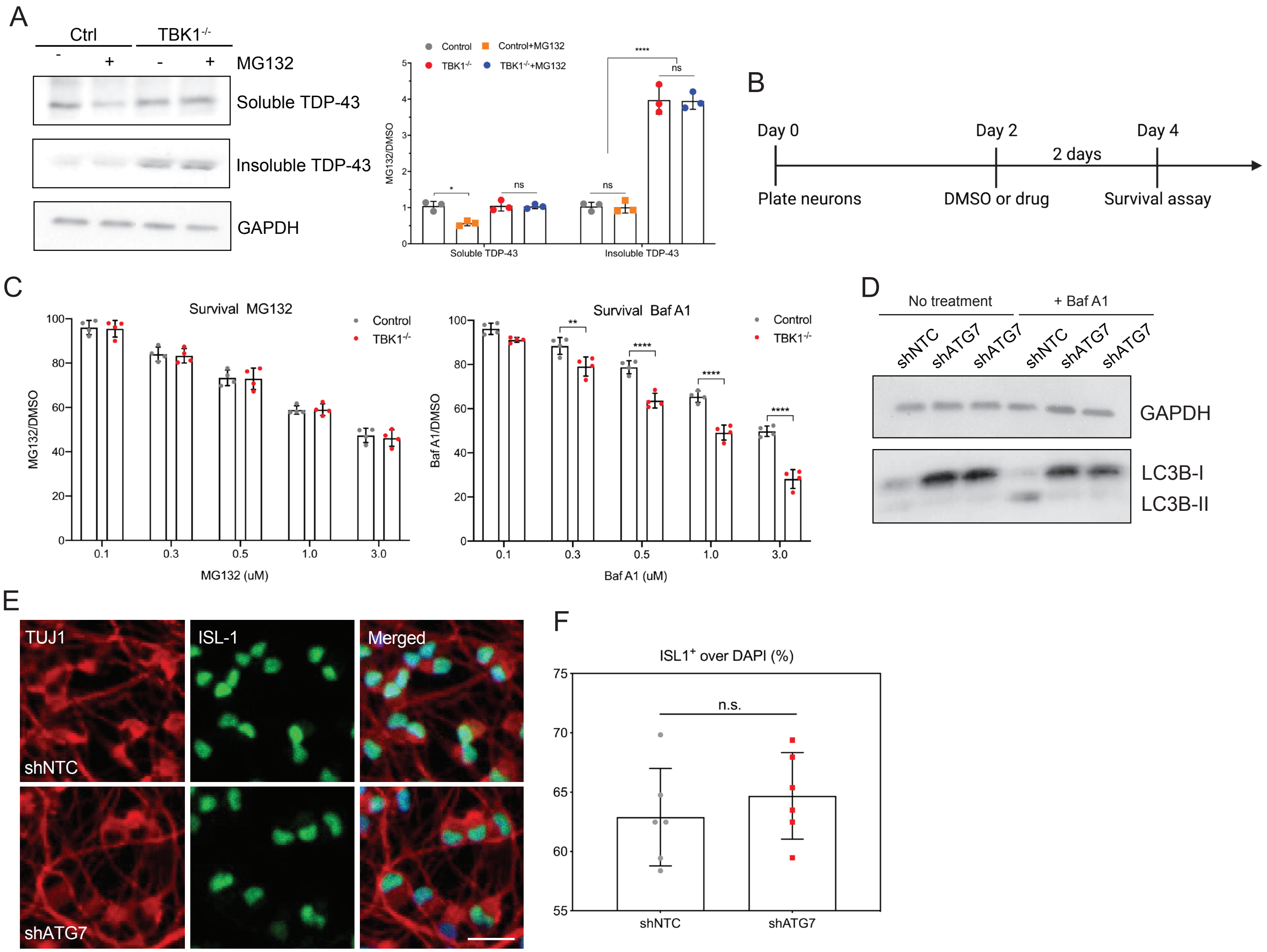

### Supplemental Figure 4

#### Supplemental Figure 4

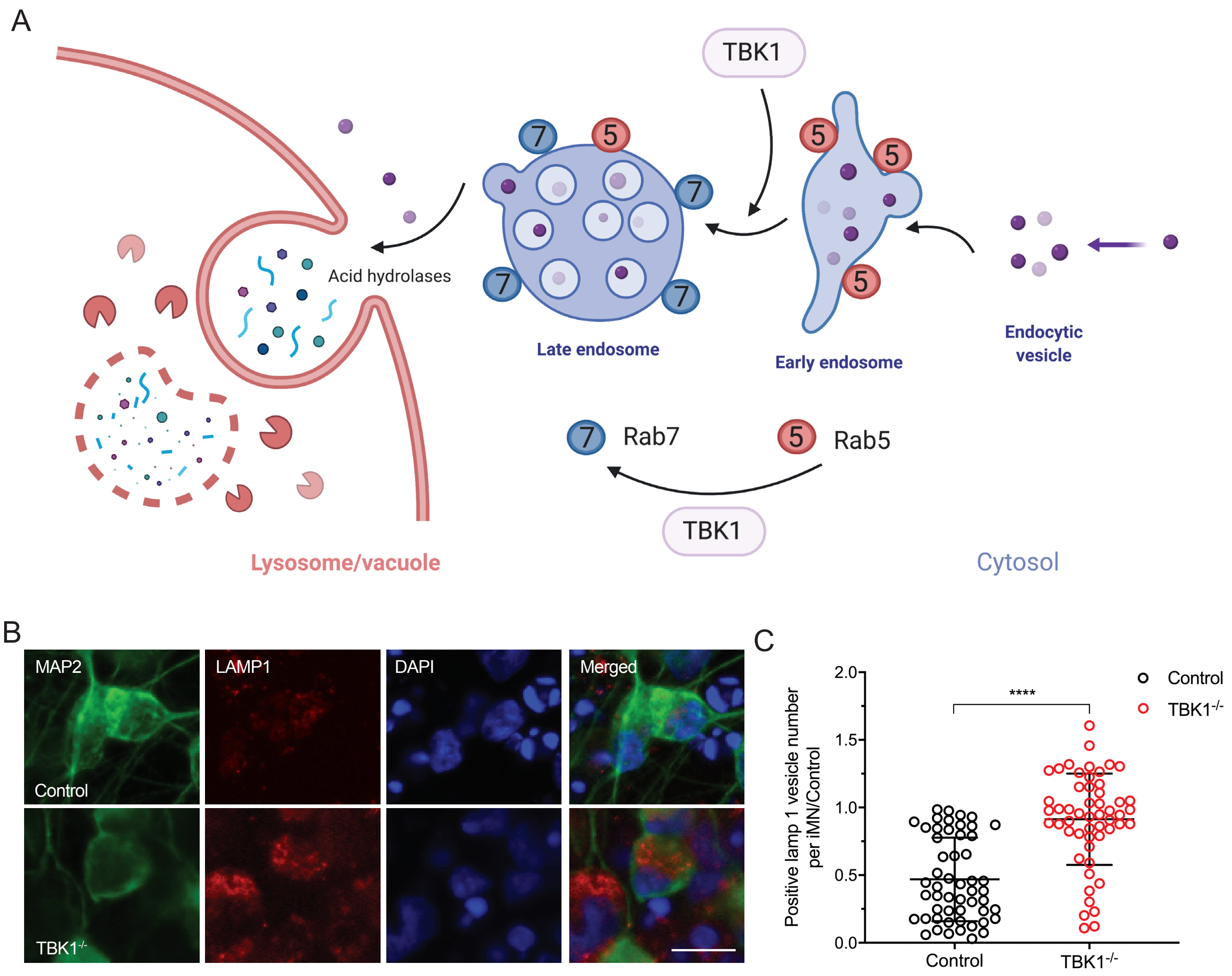

### Supplemental Figure 5

#### Supplemental Figure 5

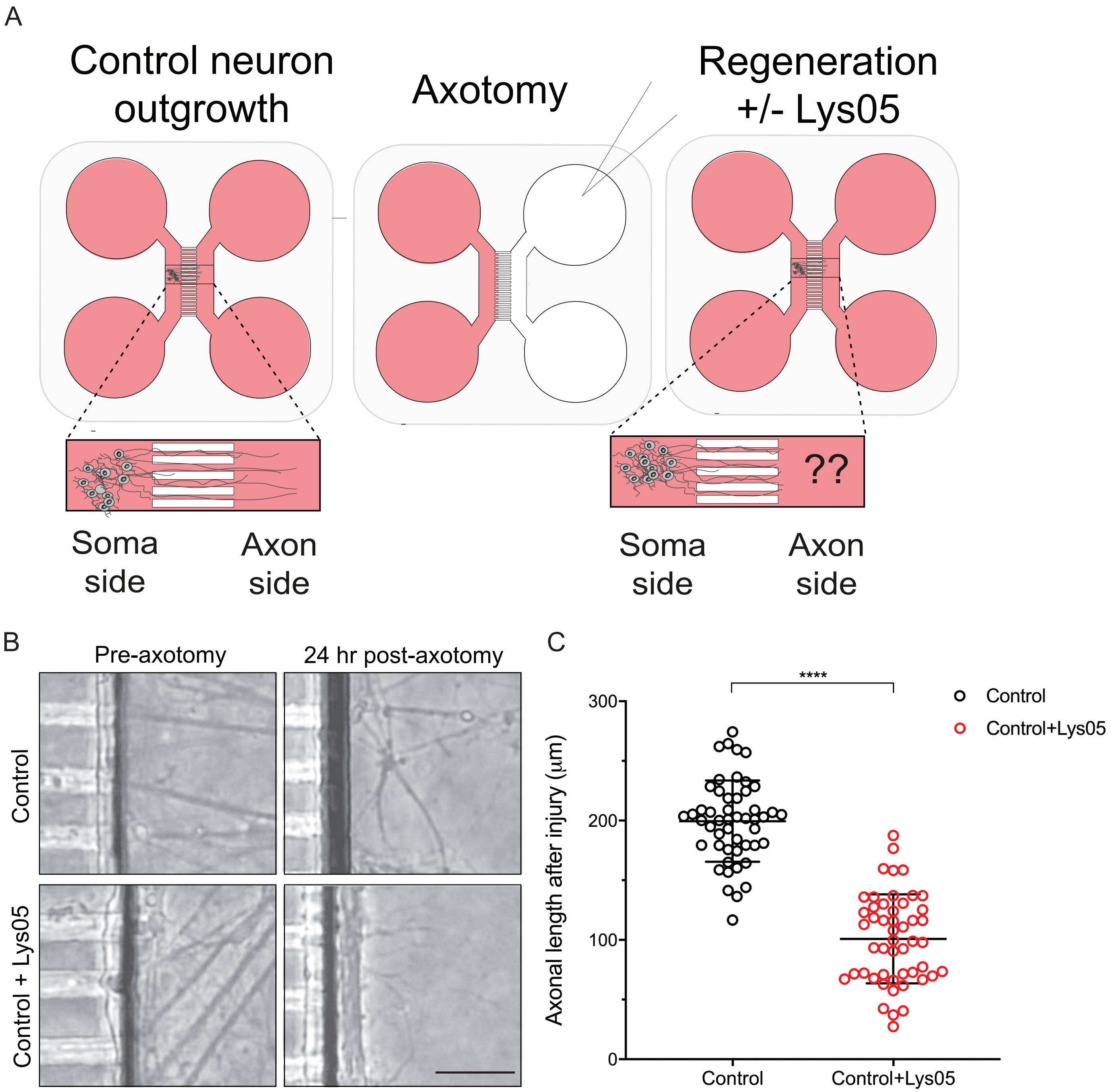

### Supplemental Figure 6

#### Supplemental Figure 6

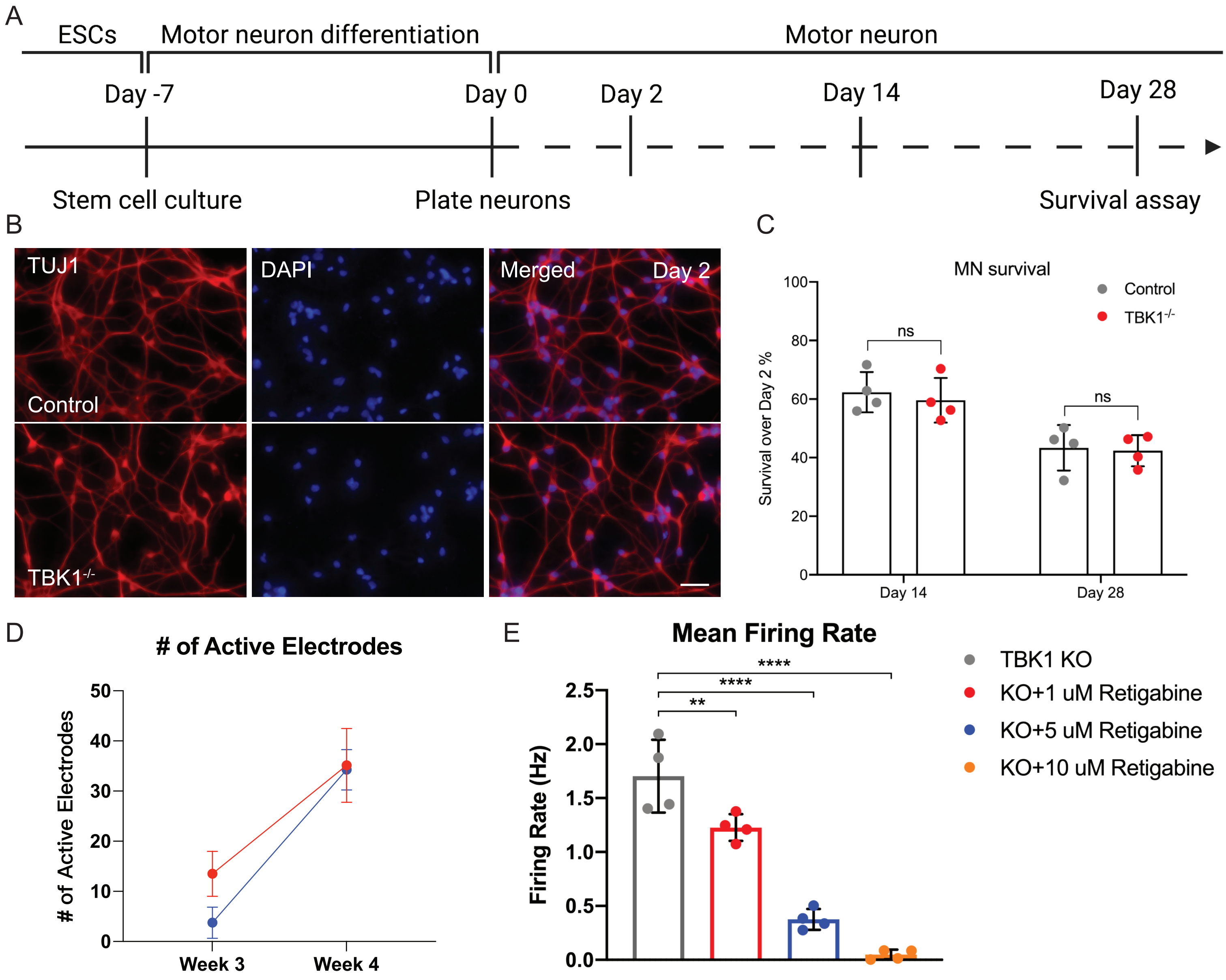

### Supplemental Figure 7

#### Supplemental Figure 7

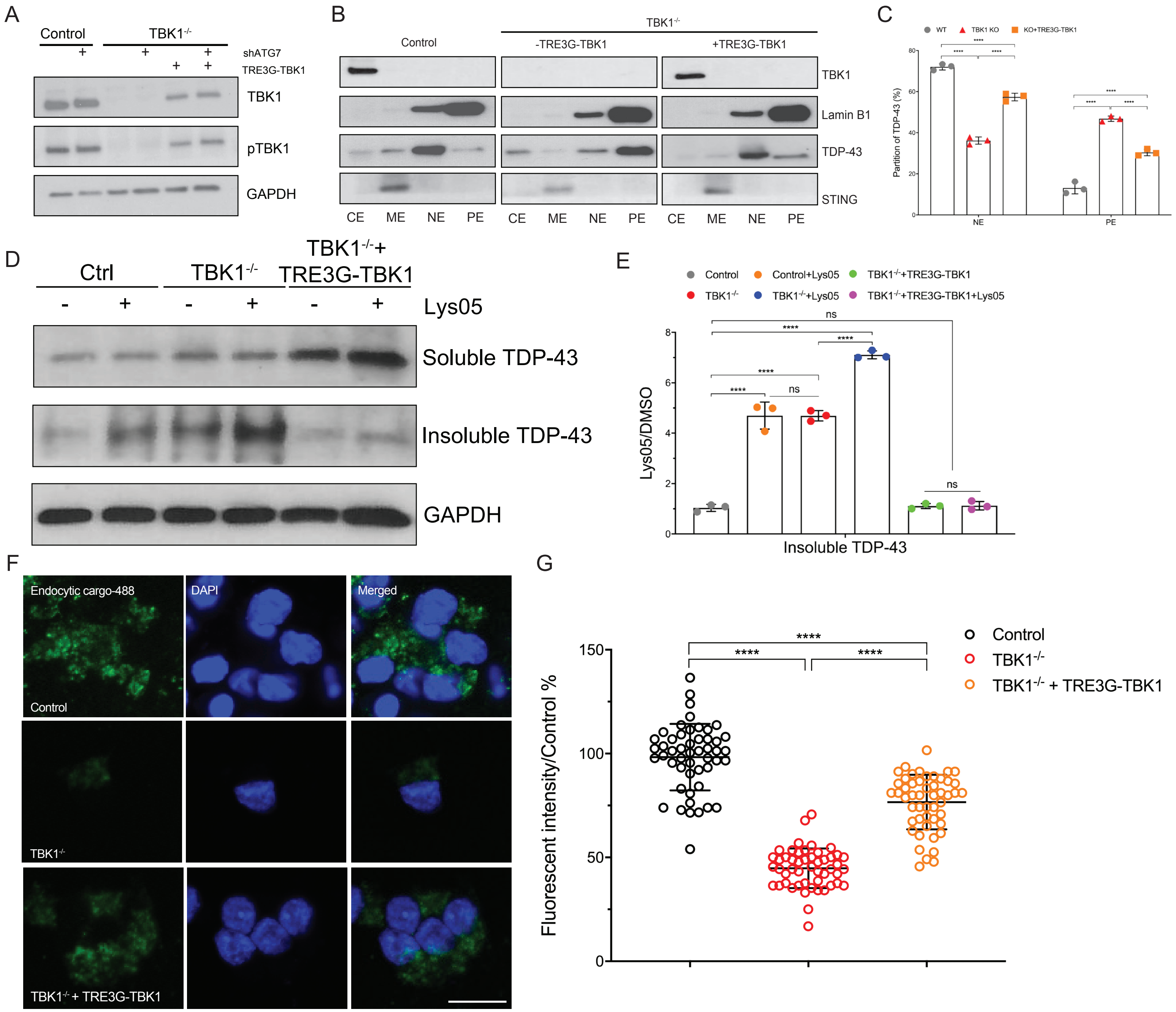

### Supplemental Figure 8

#### Supplemental Figure 8

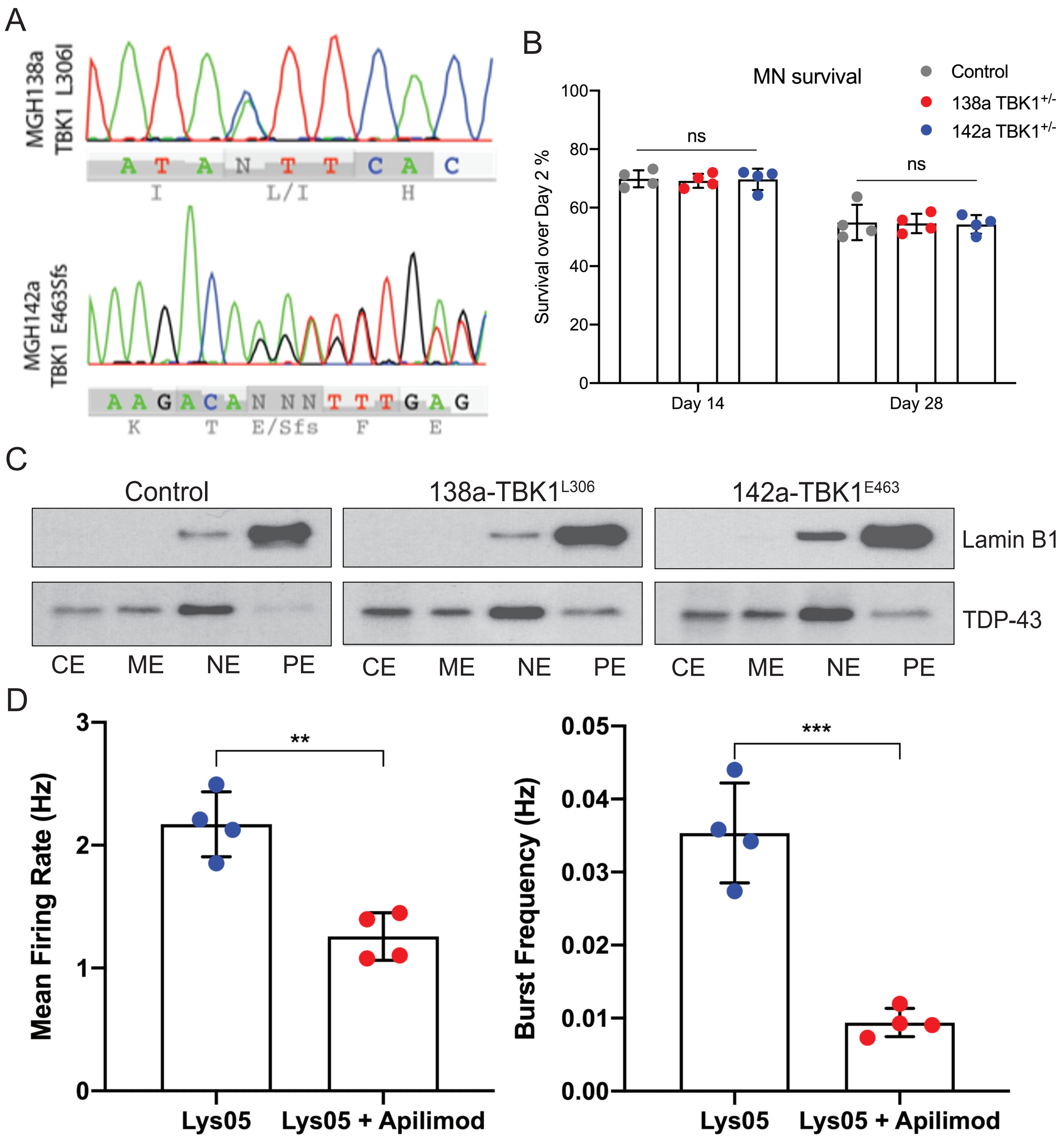
